# An engineered human cardiac tissue model reveals contributions of systemic lupus erythematosus autoantibodies to myocardial injury

**DOI:** 10.1101/2024.03.07.583787

**Authors:** Sharon Fleischer, Trevor R. Nash, Manuel A. Tamargo, Roberta I. Lock, Gabriela Venturini, Margaretha Morsink, Vanessa Li, Morgan J. Lamberti, Pamela L. Graney, Martin Liberman, Youngbin Kim, Richard Z. Zhuang, Jaron Whitehead, Richard A. Friedman, Rajesh K. Soni, Jonathan G. Seidman, Christine E. Seidman, Laura Geraldino-Pardilla, Robert Winchester, Gordana Vunjak-Novakovic

## Abstract

Systemic lupus erythematosus (SLE) is a highly heterogenous autoimmune disease that affects multiple organs, including the heart. The mechanisms by which myocardial injury develops in SLE, however, remain poorly understood. Here we engineered human cardiac tissues and cultured them with IgG fractions containing autoantibodies from SLE patients with and without myocardial involvement. We observed unique binding patterns of IgG from two patient subgroups: (i) patients with severe myocardial inflammation exhibited enhanced binding to apoptotic cells within cardiac tissues subjected to stress, and (ii) patients with systolic dysfunction exhibited enhanced binding to the surfaces of viable cardiomyocytes. Functional assays and RNA sequencing (RNA-seq) revealed that IgGs from patients with systolic dysfunction exerted direct effects on engineered tissues in the absence of immune cells, altering tissue cellular composition, respiration and calcium handling. Autoantibody target characterization by phage immunoprecipitation sequencing (PhIP-seq) confirmed distinctive IgG profiles between patient subgroups. By coupling IgG profiling with cell surface protein analyses, we identified four pathogenic autoantibody candidates that may directly alter the function of cells within the myocardium. Taken together, these observations provide insights into the cellular processes of myocardial injury in SLE that have the potential to improve patient risk stratification and inform the development of novel therapeutic strategies.

## INTRODUCTION

Myocardial involvement occurs in 25% - 50% of SLE patients, including those without symptoms, and can result in myocardial inflammation with ventricular dysfunction, progression to heart failure, and high rates of mortality^1–7^. Organ damage in SLE is presumed to be driven by specific autoantibody populations that react with host tissue, form immune complexes, and drive inflammation^8–10^. In the heart, the mechanisms by which autoantibodies contribute to myocardial injury remain elusive. Growing evidence suggests that autoantibodies can directly mediate myocardial injury. Congenital heart block in neonates exposed to maternal SLE autoantibodies through passive transfer across the placenta provides evidence that injury can occur through the direct effects of autoantibody binding^11–15^. This paradigm is further supported by evidence that autoantibodies binding to cardiac surface proteins including ion channels and β-adrenergic receptors impair calcium handling, action potential propagation, increase reactive oxygen species (ROS), activate apoptosis, and cause heart failure and arrhythmias^16–24^. In addition, multiple studies have reported direct contributions of SLE autoantibodies to alterations in brain function by modulating cellular process in neurons^25,26^. Taken together, these studies further suggest that in addition to complement- and immune cell-mediated myocardial injury, autoantibodies may directly lead to cardiac dysfunction via independent mechanisms.

Cardiomyocytes derived from human induced pluripotent stem cells (hiPSC-CMs) represent a promising approach for modeling human cardiac pathophysiology *in vitro*. We previously reported improved maturation of hiPSC-CMs within engineered human cardiac tissues by an electromechanical stimulation regimen that gradually increases the stimulation frequency (2 Hz to 6 Hz)^27^. This “intensity training” regimen subjects tissues to stress while also promoting the development of features similar to those of the adult human myocardium, including metabolism, ultrastructure and electrophysiology^27,28^.

Notably, cellular stress in SLE has been associated with the redistribution and clustering of intracellular autoantigens to apoptotic blebs on the surface of the plasma membrane^29,30^. Antigenic fragments created in these blebs, often in relation to the generation of reactive oxygen species (ROS), evoke autoimmune inflammatory responses in some SLE patients unlike those without such cellular stress^29,30^. Capitalizing on these observations, we hypothesized that exposing tissues to electromechanical stress could simulate cellular processes that exacerbate myocardial injury in SLE and thereby improve the predictive power of our model.

Here, we cultured engineered human cardiac tissue models with purified serum-derived IgG fractions from individual SLE patients to capture their specific autoantibody repertoire and effect on cardiac tissue performance. SLE patients with and without clinical evidence of myocardial inflammation (Myo+ and Myo-groups, respectively) were studied. The Myo+ group included SLE patients with new-onset systolic dysfunction (SD), defined as reduced left ventricular ejection fraction (EF) at the time of serum collection (denoted Myo+SD+), and patients with preserved EF (denoted Myo+SD-). We quantified IgG binding to cardiac tissues, and assessed their impact on contractile function, and molecular and transcriptional profiles. Using PhIP-Seq, we characterized patient-specific autoantibody targets, and with liquid chromatography tandem mass spectrometry (LC-MS/MS) we profiled cell surface accessible antigens on cardiomyocytes and cardiac fibroblasts to inform pathogenic autoantibody candidates (**Fig.1**).

**Figure 1:**
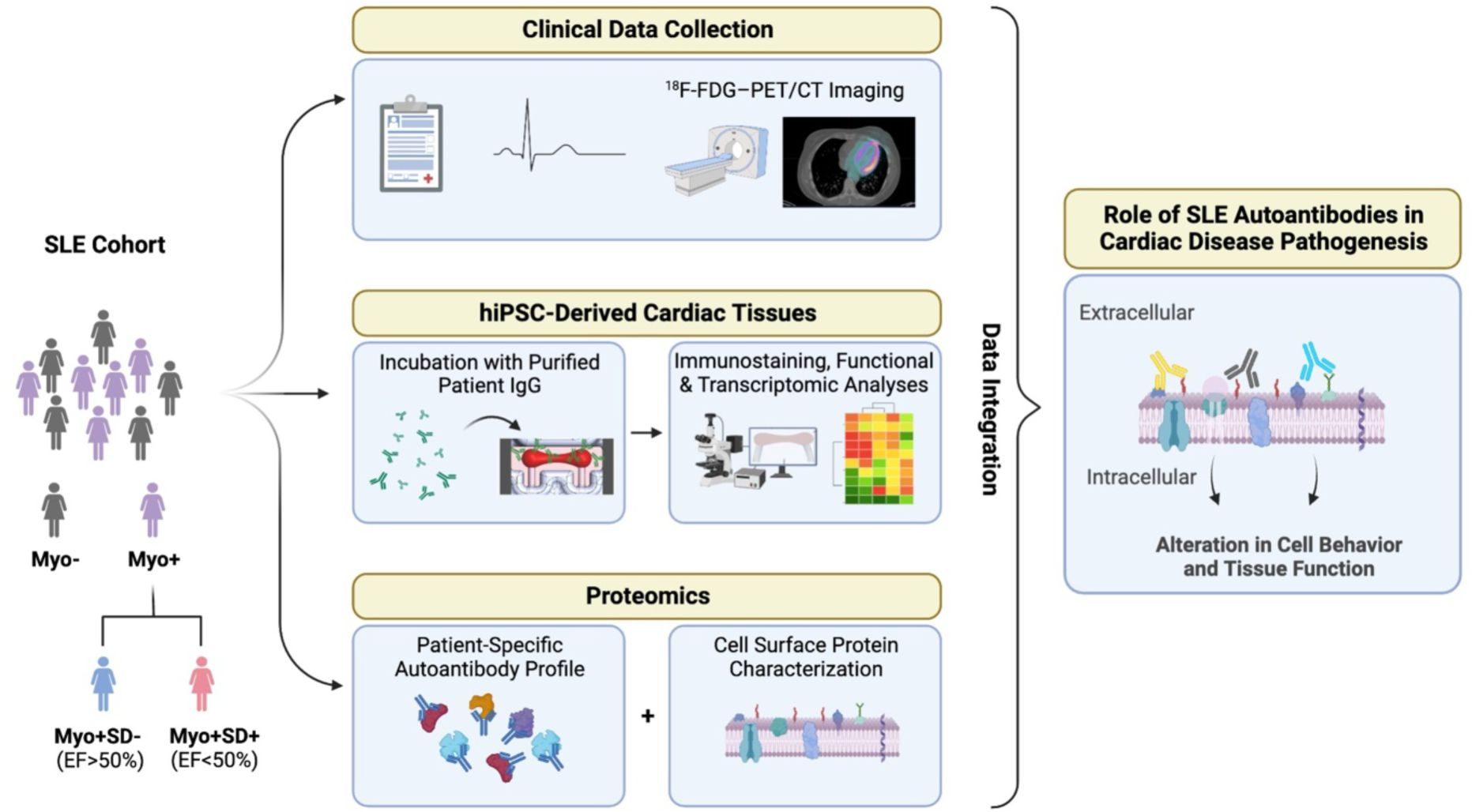
Overview of the experimental design.

## RESULTS

### SLE cohorts

Samples were obtained from patients enrolled in two SLE cohorts, one pre-existing cohort of patients without cardiac symptoms and one prospective cohort of patients hospitalized for active cardiac disease. Only patients without prior histories of cardiovascular disease or myocarditis were included. SLE was diagnosed using the 1997 American College of Rheumatology classification criteria^31^ inclusive of medical history and exam, and clinical studies with ^18^F -fluorodeoxyglucose uptake in positron emission tomography/computed tomography (^18^F-FDG–PET/CT) were conducted to determine the presence or absence of myocardial inflammation. All ^18^F-FDG–PET/CT scans were evaluated qualitatively for the presence or absence of enhanced myocardial ^18^F-FDG uptake, and levels of such uptake were quantified by standardized uptake values (SUV)^32^. SLE patients were divided into two primary groups based on myocardial ^18^F-FDG uptake: (i) Myo-patients were without evidence of myocardial inflammation (SUVmax < 1.5; n=3) and (ii) Myo+ patients had evidence of myocardial inflammation (SUVmax ≥ 1.5; n=8). At the time of ^18^F-FDG–PET/CT, serum samples were collected, and echocardiograms obtained to identify Myo+ patients with systolic dysfunction (Myo+SD+, n=4, with left ventricular ejection fraction (EF) < 50%) and with preserved systolic function (Myo+SD-, n=4, EF ≥ 50%). No patients in either group had elevated serum troponin, and no Myo+SD+ patients had histories of reduced EF prior to these studies, suggesting recent onset systolic dysfunction. Detailed clinical data are summarized in **Table 1**.

**Table 1:** Patient cohort clinical characteristics. All patients in the cohort have SLE. Patients were grouped based on the presence or absence of myocardial inflammation (Myo+ and Myo-, respecitively); the patients with myocardial inflammation were further sub-grouped based on systolic dysfunction (SD). Myocardial ^18^F-FDG uptake was not quantified in all patients and is reported as positive or negative by qualitative analysis. Abbreviations: SUV, standardized uptake values; EF, left ventricular ejection fraction; ACR, American College of Rheumatology; SLEDAI, systemic lupus erythematosus disease activity index; QTc, corrected QT interval; ESR, erythrocyte sedimentation rate; CRP, c-reactive protein; B2GP, beta-2-glycoprotein; ACL, anticardiolipin.

| Patient Descriptor | Myo- 1 | Myo- 2 | Myo- 3 | Myo+SD- 1 | Myo+SD- 2 | Myo+SD- 3 | Myo+SD- 4 | Myo+SD+ 1 | Myo+SD+ 2 | Myo+SD+ 3 | Myo+SD+ 4 |
| --- | --- | --- | --- | --- | --- | --- | --- | --- | --- | --- | --- |
| Group | Myocardial Inflammation (-) |  |  | Myocardial Inflammation (+) |  |  |  |  |  |  |  |
| Sub Group | N/A |  |  | Systolic Dysfunction (-) |  |  |  | Systolic Dysfunction (+) |  |  |  |
| Age | 30 | 45 | 26 | 37 | 38 | 34 | 25 | 23 | 43 | 46 | 39 |
| Sex | Female | Female | Female | Male | Female | Female | Female | Female | Female | Female | Female |
| Disease Duration (years) | 10 | 34 | 5 | 8 | 4 | 12 | 8 | 5 | 7 | 22 | 28 |
| EF (%) | 55-60 | 60-65 | 62 | 55 | 60-65 | 60-65 | 58 | 35 | 47 | 45-50 | 10-15 |
| ACR Criteria | 4 | 10 | 7 | 10 | 3 | 7 | 9 | 11 | 11 | 11 | 5 |
| SLEDAI | 4 | 0 | 4 | 2 | 4 | 5 | 5 | 18 | 10 | 0 | 0 |
| QTc (ms) | 423 | 428 | 467 | 416 | 438 | 437 | 411 | 493 | 486 | 438 | 488 |
| Troponin (ng/mL) | <0.01 | <0.01 | <0.01 | <0.01 | <0.01 | <0.01 | <0.01 | <0.01 | <0.01 | <0.01 | <0.01 |
| ESR (mm/hr) | 17 | 27 | 44 | 28 | 22 | 45 | 5 | 57 | 79 | 55 | 60 |
| CRP (mg/L) | 3.2 | 1.41 | 5.37 | 0.25 | 2 | 0.84 | 0.3 | 12.4 | 5.88 | 1.65 | 67.27 |
| C3 (mg/dL) | 126 | 61 | 125 | 92 | 133 | 121 | 77 | 144 | 119 | 83 | 163 |
| C4 (mg/dL) | 28 | 5 | 36 | 30 | 33 | 24 | 8 | 26 | 22 | 20 | 23 |
| anti-dsDNA Antibody | <6 | 245.5 | 47.9 | 121.2 | 14.9 | 7.6 | 13.5 | 52.2 | 62.1 | 24 | 7.4 |
| SSA Antibody | 0.9 | <0.2 | <0.2 | >8 | 3.7 | >8 | <0.2 | >8 | <0.2 | <0.2 | <0.2 |
| SSB Antibody | <0.2 | <0.2 | <0.2 | >8 | <0.2 | 0.5 | <0.2 | >8 | <0.2 | <0.2 | <0.2 |
| Smith Antibody | <0.2 | 0.9 | <0.2 | 0.6 | <0.2 | <0.2 | <0.2 | 7.3 | 0.3 | <0.2 | <0.2 |
| B2GP IgG | 0 | 0 | 0 | 0 | 0 | 0 | 1 | 1 | 2 | 17 | 13 |
| B2GP IgM | 3 | 1 | 0 | 2 | 2 | 2 | 7 | 2 | 5 | 0 | 34 |
| ACL IgG | 5 | 2 | 2 | 0 | 2 | 4 | 2 | 0 | 1 | 18 | 28 |
| ACL IgM | 2 | 5 | 0 | 0 | 0 | 3 | 10 | 0 | 5 | 0 | 11 |

### Engineered human cardiac tissue with advanced functional and metabolic properties

Cardiac tissues were engineered by resuspending hiPSC-CMs and primary human cardiac fibroblasts at a 3:1 ratio in fibrin hydrogel stretched between two flexible pillars, and were electromechanically stimulated using the milliPillar platform^28^. Electromechanical stimulation frequency was increased from 2 to 6 Hz over two weeks, followed by an additional week of stimulation at 2 Hz^19^. To further enhance tissue performance, tissues were cultured in maturation medium with substrates that shifted cardiomyocyte metabolism from glycolysis toward fatty acid oxidation, increased mitochondrial content and optimized function^33^ (detailed in Methods; **Fig. 2a**, top panel). Tissues exhibited enhanced compaction, α-actinin striations, and calcium handling as compared to those cultured in standard medium (**Extended Data Fig. 1a-d**). Tissue stimulation at 6 Hz caused significantly increased release of lactate dehydrogenase (LDH), indicative of cellular stress (**Extended Data Fig. 1e**).

**Figure 2:**
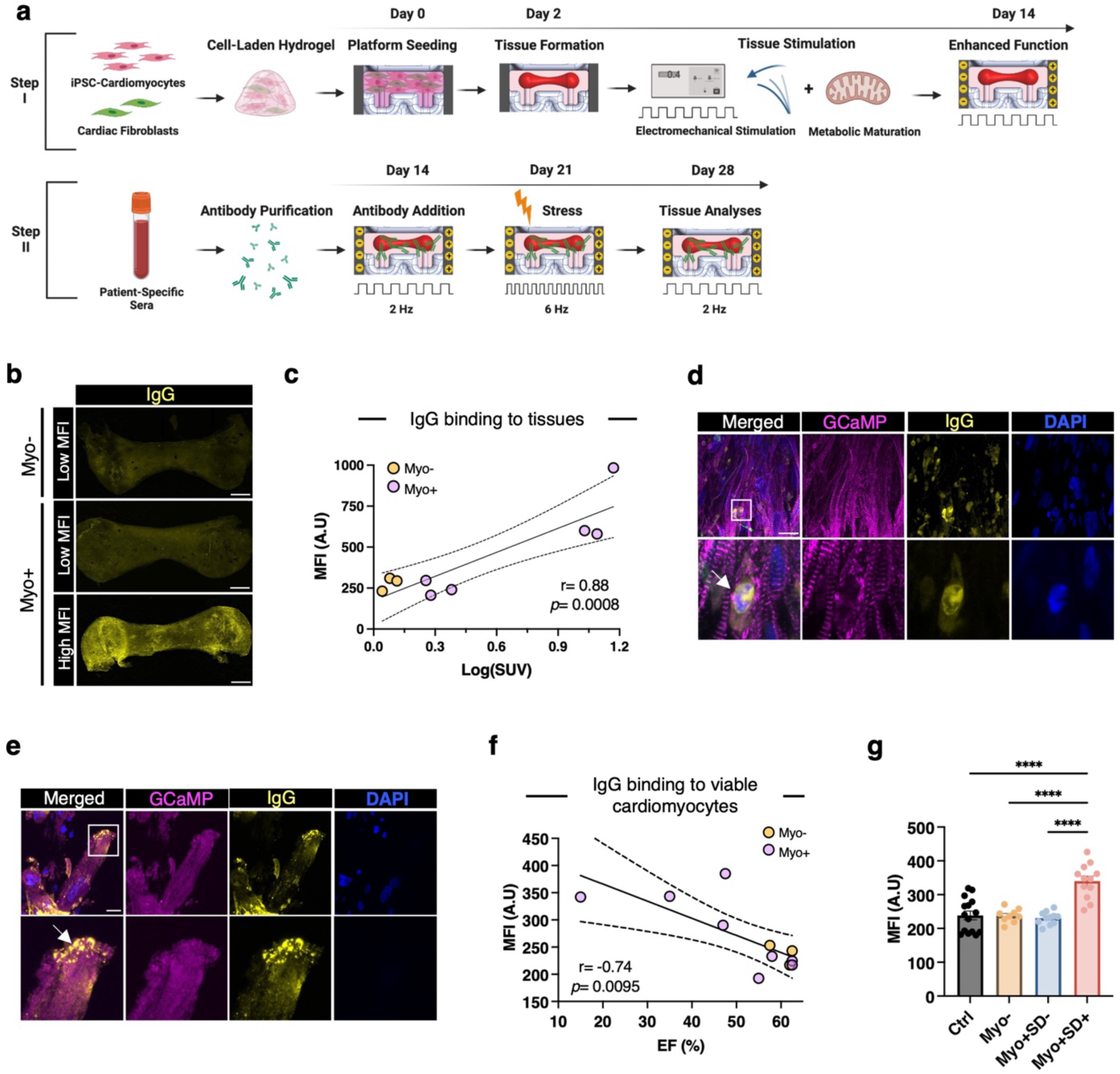
Distinct patient IgG reactivity with engineered cardiac tissues and hiPSC-cardiomyocytes corresponded with specific clinical outcomes. (**a**) Overview of tissue formation, maturation, and treatment with purified patient-derived IgG. (**b**) Indirect immunofluorescence staining of human IgG (yellow) bound to engineered cardiac tissues (scale bar: 500 μm). (**c**) Log-linear correlation between clinical measurements of myocardial inflammation (^18^F-FDG uptake quantified as SUVs) and IgG binding levels to engineered cardiac tissues (MFI). Each dot represents a patient (n = 9). Correlation strength measured by Pearson’s *r*. (**d,e**) Immunofluorescence staining of engineered cardiac tissues showing strong IgG binding to condensed nuclei (**d**, scale bar: 20 μm, bottom panels are higher magnification of region of interest in top panels, top panel is confocal maximum intensity projection, bottom panel is single confocal plane) and apoptotic blebs on the cell surface (**e**, scale bar: 20 μm, bottom panels are higher magnification of region of interest in top panels, both panels are single confocal planes). (**f**) Linear correlation between patient EF and corresponding IgG binding levels to the cell surface of viable hiPSC-derived cardiomyocytes (MFI). Each dot represents a patient (n = 9). Correlation strength measured by Pearson’s *r*. (**g**) Quantification of patient IgG binding to viable hiPSC-cardiomyocytes based on clinical subgroupings. Data are the mean ± s.e.m; individual values are indicated by dots; n = 9-15 independent staining experiments per group (n = 3 technical replicates for each patient IgG sample); one-way ANOVA with Tukey’s test for multiple comparisons; **** p < 0.001.

### IgG from SLE Myo+ patients with elevated myocardial inflammation exhibited enhanced binding to stressed cardiac tissues

IgG fractions were purified and quantified from each patient’s serum samples. At day 14 of cardiac tissue culture, IgG fractions from individual patients were added and tissues were cultured for 14 additional days (**Fig. 2a**, bottom panel). The extent of serum IgG binding to tissues was evaluated by indirect immunostaining using secondary anti-human IgG and quantified as mean fluorescent intensity (MFI). All Myo-patients had low MFI levels. A subgroup of Myo+ patients exhibited low MFI levels, while other Myo+ patients had substantially higher MFI levels (**Fig. 2b, Extended Data Fig. 2**). MFI studies of an additional SLE Myo+ patient with high levels of myocardial FDG uptake (SUVmax = 12.4) confirmed this trend. Linear regression analysis (inclusive of all patients) revealed a significant positive correlation between MFI and the levels of myocardial inflammation, as determined by ^18^F-FDG–PET/CT (r= 0.88, **Fig. 2c**). MFI correlated more strongly with myocardial inflammation than did any clinical metric of myocardial health or SLE severity (**Extended Data Fig. 3a**).

To further probe the cellular targets of IgGs, tissues were co-stained with secondary anti-human IgG and α-actinin to evaluate cardiomyocyte integrity and IgG localization. The extent of IgG binding within the stressed tissues varied from cell to cell (**Fig. 2d**). The subset of cells with the greatest IgG staining were cardiomyocytes with disrupted α-actinin striations and condensed nuclei, with IgG binding primarily to the nuclei, indicating late cell apoptosis with loss of cell membrane integrity that allowed IgG penetration (**Fig. 2d)**. Additionally, we identified another subset of cells with high levels of localized IgG binding to presumptive apoptotic blebs on surfaces of cardiomyocytes with minimal nuclear condensation (**Fig. 2e**).

### IgG from SLE Myo+SD+ patients exhibit enhanced binding to viable cardiomyocytes

As previous studies report that autoantibodies bound to plasma membrane targets can directly exert biological effects^11,16–20^, we investigated the patterns of IgG binding to the surfaces of the viable cardiomyocyte populations. iPSC-CMs were incubated with patient IgGs, followed by incubation with secondary anti-human IgG, and sorted using flow cytometry to exclude dead and apoptotic cells (**Methods**; **Extended Data Fig. 3b**). MFI levels of IgG binding to the surface of viable cardiomyocytes did not correlate with clinical assessments of myocardial inflammation (**Extended Data Fig. 3c**). However, a significant negative correlation with EF was observed (r = −.74, **Fig. 2f**). Indeed, Myo+SD+ SLE patients exhibited significantly higher binding levels of IgG to viable cardiomyocytes compared to all other patient groups and healthy controls (**Fig. 2g**).

### IgG from Myo+SD+ patients altered engineered cardiac tissue cellular composition and function

We next sought to investigate whether IgG binding to the surface of viable iPSC-CMs exerted functional effects. After 14 days of incubation with IgGs from individual SLE patients or healthy controls (**Fig. 3a**) tissues were studied by immunofluorescence staining for α-actinin and vimentin. We observed a strong negative correlation (r= −.8) between the levels of vimentin staining, indicative of fibroblast content, and patient EFs (**Fig. 3b,c**).To confirm this trend, we treated fibroblasts in monolayers with IgG from Myo+SD+ patients and observed a significant increase in Ki-67 staining, indicating increased fibroblast proliferation (**Fig. 3d**).

**Figure 3:**
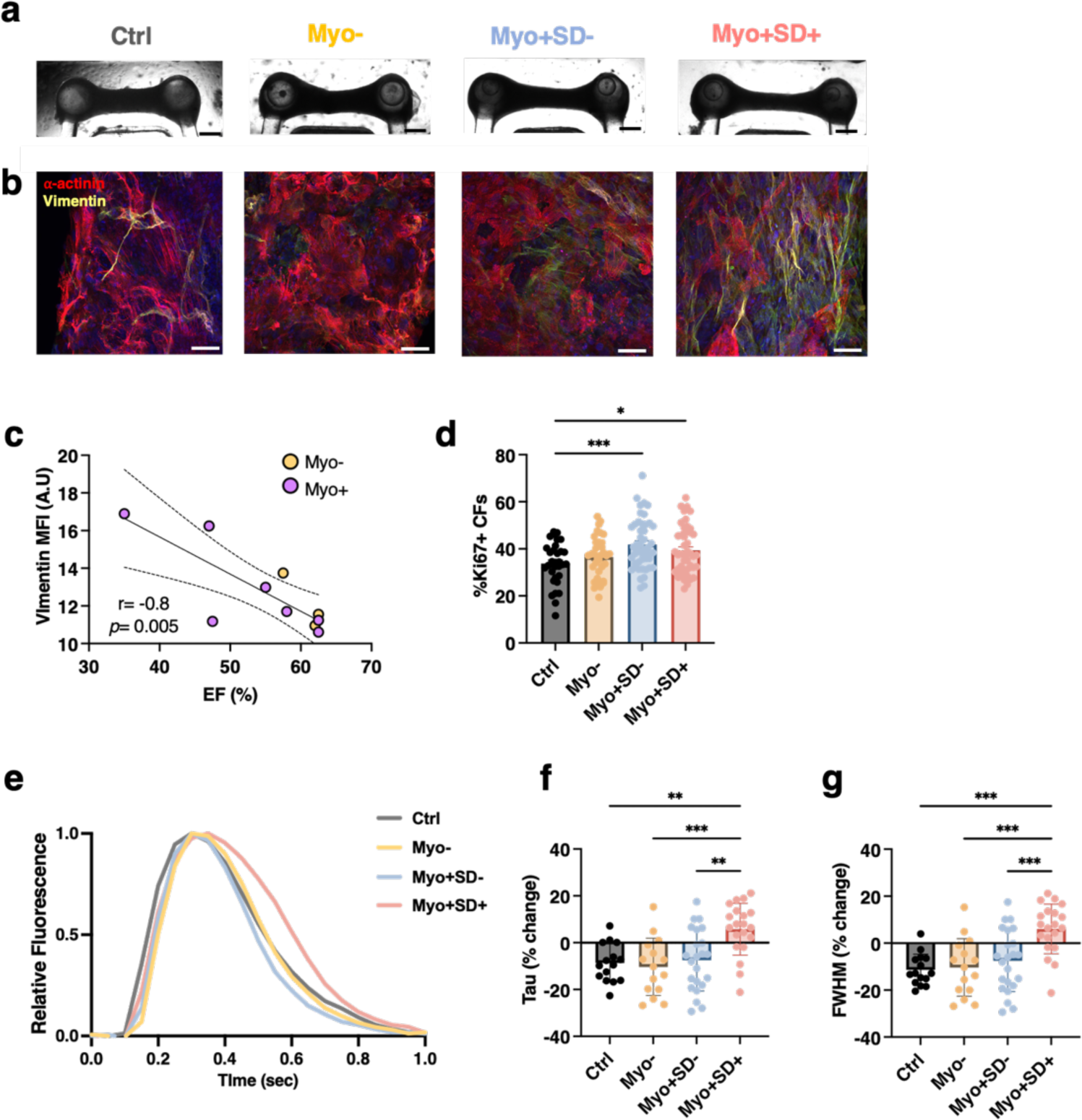
IgG from SLE patients with myocardial inflammation and systolic dysfunction (Myo+SD+) altered engineered cardiac tissue function and composition. (**a**) Brightfield images of engineered cardiac tissue following treatment with patient IgG showing overall tissue morphology (scale bar: 500 μm). (**b**) Representative immunofluorescence images of engineered cardiac tissue stained to show cardiomyocytes (α-actinin, red) and fibroblasts (vimentin, yellow) following treatment with IgG from patients in each subgroup (scale bar: 50 μm). (**c**) Linear correlation between patient ejection fraction (EF) and vimentin staining intensity of tissues treated with the corresponding patient IgG. Each dot represents a patient (n = 11). Correlation strength measured by Pearson’s *r*. (**d**) Quantification of the percentage of Ki67+ cardiac fibroblasts following treatment with patient IgG. Data are the mean ± s.e.m; individual values are indicated by dots; n= 33-48 independent staining experiments per group (n = 11-12 technical replicates for each patient IgG sample); ordinary one-way ANOVA with Dunnett’s test for multiple comparisons; * p < 0.05, *** p < 0.001. (**e**) Representative traces of calcium flux in engineered cardiac tissues following treatment with patient IgG. (**f, g**) Quantification of the parameters tau (f) and FWHM (g) extracted from engineered cardiac tissue calcium transients following treatment with patient IgG. Data are the mean ± s.e.m; individual values are indicated by dots; n = 15-22 independent engineered tissues per group (n = 5-6 technical replicates for each patient IgG sample); ordinary one-way ANOVA with Tukey’s test for multiple comparisons; * p < 0.05, ** p < 0.005, *** p < 0.001. MFI, mean fluorescent intensity; CFs, cardiac fibroblasts; FWHM, full width half max.

As autoantibodies can alter cardiomyocytes calcium homeostasis^17,20^, we assessed cardiac tissue contractile function and calcium handling using non-invasive data acquisition and analysis modalities (**Extended Data Fig. 4a-d**). In comparison to baseline, prior to IgG addition, calcium transient decay constant (tau, τ) and transient width (full width half max, FWHM) were significantly higher in tissues cultured with IgGs from Myo+SD+ patients compared with all other groups (**Fig. 3 e-g**). These results were further supported by dose-dependent increases in tau and FWHM in hiPSC-CMs monolayers cultured with IgGs from Myo+SD+ patients (**Extended Data Fig. 5a,b**). There were no significant differences in force generation.

### The binding of IgG from Myo+SD+ patients to cardiac tissues altered tissue transcriptomics

Bulk RNA-seq identified 707 differentially expressed genes (DEGs, FDR<0.05) between tissues incubated with IgGs from Myo+SD+ and Myo-patients, and 1,035 DEGs between tissues treated with IgGs from Myo+SD+ and Myo+SD-patients (**Fig. 4a,b, Supplementary Information Table 1)**. No DEGs were found between tissues treated with IgGs from Myo+SD- and Myo-patients, or between tissues treated with IgGs from Myo-patients and healthy controls (**Fig. 4a, Extended Data Fig. 6a**).

**Figure 4:**
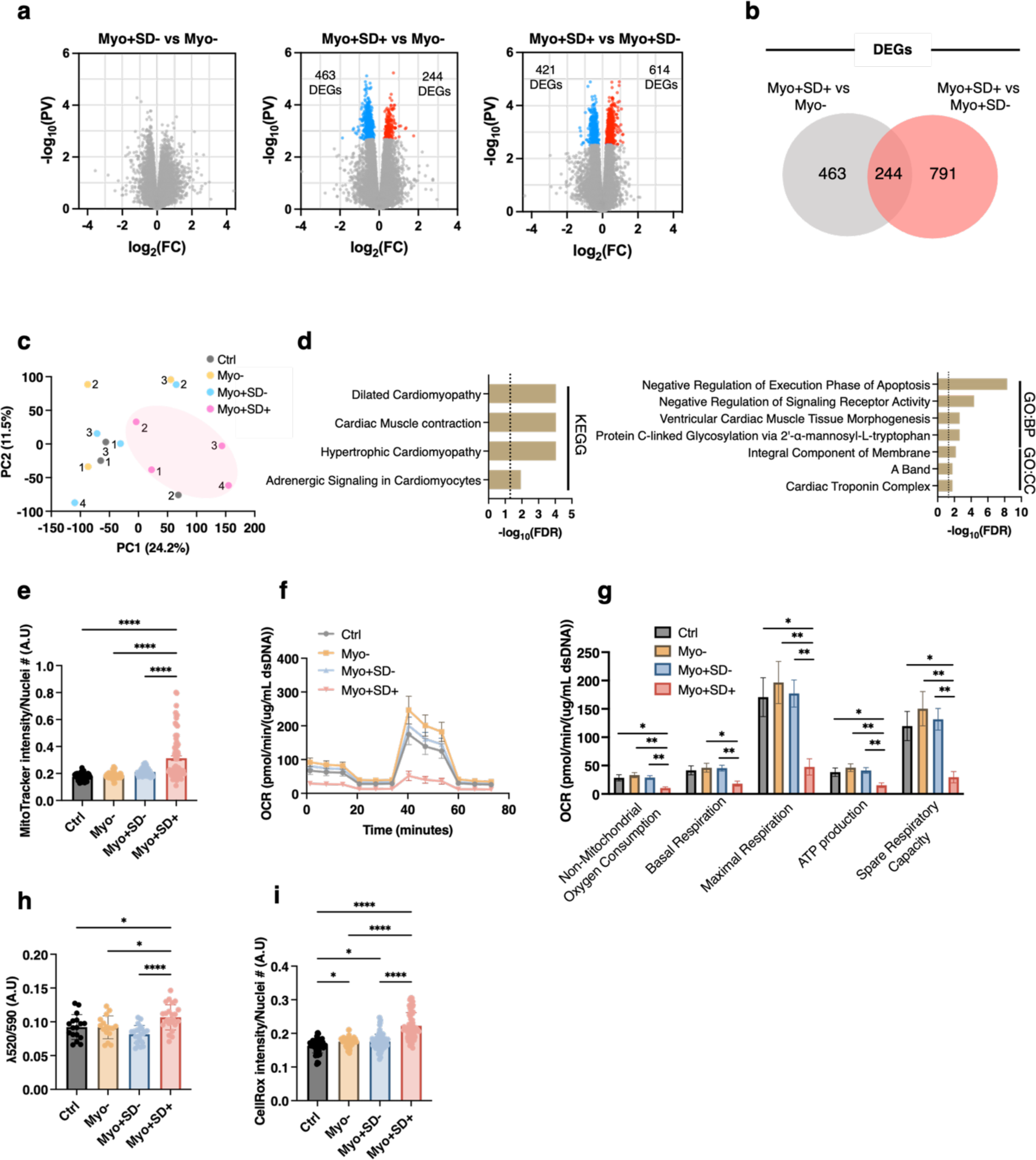
IgG isolated from SLE patients with myocardial inflammation and systolic dysfunction (Myo+SD+) led to differential tissue transcriptomics and impaired mitochondrial function. (**a**) Volcano plots depicting the differential expression between tissues cultured with IgG from the different clinical groups. Genes that are significantly upregulated (FDR < 0.05) are shown in red and genes that are significantly downregulated (FDR < 0.05) are shown in blue. (**b**) Venn diagram describing the number of DEGs between tissues treated with IgG from Myo+SD+ and either SLE patients with no myocardial inflammation (Myo-) or patients with myocardial inflammation and no systolic dysfunction (Myo+SD-). (**c**) PCA plot of global gene expression profiles for tissues treated with IgG from different patients. Each point represents the average expression of n=3 independent tissues treated with IgG from the same patient. Numbers refer to patient IDs within each clinical group. (**d**) KEGG, GO biological process (GO:BP), and GO cellular compartment (GO:CC) pathway analysis of DEGs between the Myo+SD+ and the Myo+SD-groups. (**e**) Mitochondrial abundance in hiPSC-cardiomyocytes treated with patient IgG from the different groups as measured by MitoTracker fluorescence normalized to the number of nuclei. n = 67-87 fields of view from (n = 6 independent technical replicates for each patient IgG sample; n = 10-16 fields of view per replicate); ordinary one-way ANOVA with Tukey’s test for multiple comparisons; **** p < 0.0001. (**f, g**) Characterization of mitochondrial performance by Seahorse metabolic flux assay. Data are the mean ± s.e.m; individual values are indicated by dots. n = 8-16 independent assays per group (n = 3-4 technical replicates for each patient IgG sample); ordinary one-way ANOVA with Tukey’s test for multiple comparisons; * p < 0.05, ** p < 0.01. (**h**) Changes in mitochondrial membrane potential measured by the JC-10 probe. Increasing 520/590 ratios indicate increasing membrane depolarization and mitochondrial dysfunction. Data are the mean ± s.e.m; individual values are indicated by dots. n = 16-24 independent assays per group (n = 5-6 technical replicates for each patient IgG sample); ordinary one-way ANOVA with Tukey’s test for multiple comparisons; * p < 0.05, **** p < 0.0001. (**i**) ROS accumulation as measured by CellRox fluorescence normalized to the number of nuclei. Data are the mean ± s.e.m; individual values are indicated by dots. n = 65-86 fields of view from (n = 6 independent technical replicates for each patient IgG sample; n = 10-16 fields of view per replicate); ordinary one-way ANOVA with Tukey’s test for multiple comparisons; * p < 0.05, **** p < 0.0001. DEGs, differentially expressed genes; PCA, principal component analysis; OCR, oxygen consumption rate.

DEGs in tissues incubated with IgGs from Myo+SD+ patients included reduced expression of voltage-gated calcium channel subunits (*CACNB2*, *CACNA2D1*), a finding consistent with the observed impairment in calcium handling. Genes involved in aerobic respiration (*NDUFB11*, *UQCC3*, *COX6B1*) and oxidative stress (*BAD*, *ROMO1*, *SOD2*) were upregulated, and the expression of cardiomyocyte transcription factors (*MEF2A*, *MEF2C*) was decreased (**Extended Data Fig. 7a-d, Supplementary Information Table 1**).

PCA and hierarchical clustering analysis of gene expression of tissues incubated with IgGs delineated Myo+SD+ patients from other SLE patients and healthy controls **(Fig. 4c, Extended Data Fig. 6b)**. Kyoto Encyclopedia of Genes and Genome (KEGG) pathway analysis and gene ontology (GO) of DEGs between tissues treated with IgGs from Myo+SD+ and Myo+SD-patients revealed terms related to cardiomyopathies, cardiac muscle contraction and morphogenesis, adrenergic signaling, and apoptosis (**Fig. 4d**).

### IgGs from Myo+SD+ patients impaired mitochondrial function and increased oxidative stress in cardiomyocytes

Based on the high metabolic demand of cardiomyocytes and the association between mitochondrial impairment and cardiac dysfunction^34–36^, we investigated cardiomyocyte-specific gene expression from bulk RNA-seq data predicted by CIBERSORTx^37^ (**Supplementary Information Table 2**). Among the DEGs in the tissues treated with IgG from Myo+SD+, were genes associated with mitochondria and oxidative stress. KEGG analysis of these DEGs identified pathways associated with oxidative phosphorylation, ROS, and cardiomyopathy (**Extended Data Figure 8a,b**).

To confirm these observations, we treated monolayers of hiPSC-CMs with IgGs and assessed mitochondrial content using a fluorescent probe that specifically accumulates in the mitochondria of live cells. At 72 hours, iPSC-CMs cultured with Myo+SD+ IgGs displayed significantly higher mitochondrial content (**Fig. 4e**), consistent with the RNA-seq data. In contrast, Seahorse metabolic flux assays indicated that Myo+SD+ IgGs significantly decreased basal respiration, ATP production, and maximum respiratory capacity (**Fig. 4f,g**). We further explored whether this decrease in cellular respiration was associated with mitochondrial damage. Using the JC-10 probe to indicate mitochondrial membrane potential, we found that Myo+SD+ IgG-treated iPSC-CMs had significantly decreased membrane polarization potential (**Fig. 4h**). In addition, and in line with the RNA-seq data, we observed a significant increase in cellular ROS production (**Fig. 4i**).

### Myo+SD+ patients exhibited unique autoantibody repertoires

Next, we sought to investigate whether the observed phenotypic and clinical variation between patients was associated with unique autoantibody signatures. We utilized phage immunoprecipitation sequencing (PhIP-Seq), a high-throughput unbiased technique that incorporates phage display of synthetic peptides covering the entire human proteome to characterize autoantibody reactivities at scale^38^. We initially validated PhIP-Seq target detection by finding a linear correlation between read counts for SSA/Ro peptides, the most frequent antigens targeted by SLE antibodies, and clinical measurements of anti-SSA/Ro antibodies (**Extended Data Fig. 9a**). PhIP-Seq analyses of SLE serum demonstrated elevated levels of autoantibodies against 147 proteins compared to healthy controls, with high heterogeneity among patients (**Extended Data Fig. 9b, Supplementary Information Table 3a-d**). PCA of differentially detected antigen targets clearly delineated all SLE patients from healthy controls with tight clustering among patient subgroup (**Fig. 5a**). Thirty proteins were uniquely detected by Myo+SD+ autoantibodies, 26 of which are expressed in human heart^39^ (**Fig. 5b, Extended Data Fig. 10a, Supplementary Information Table 3e**).

**Figure 5:**
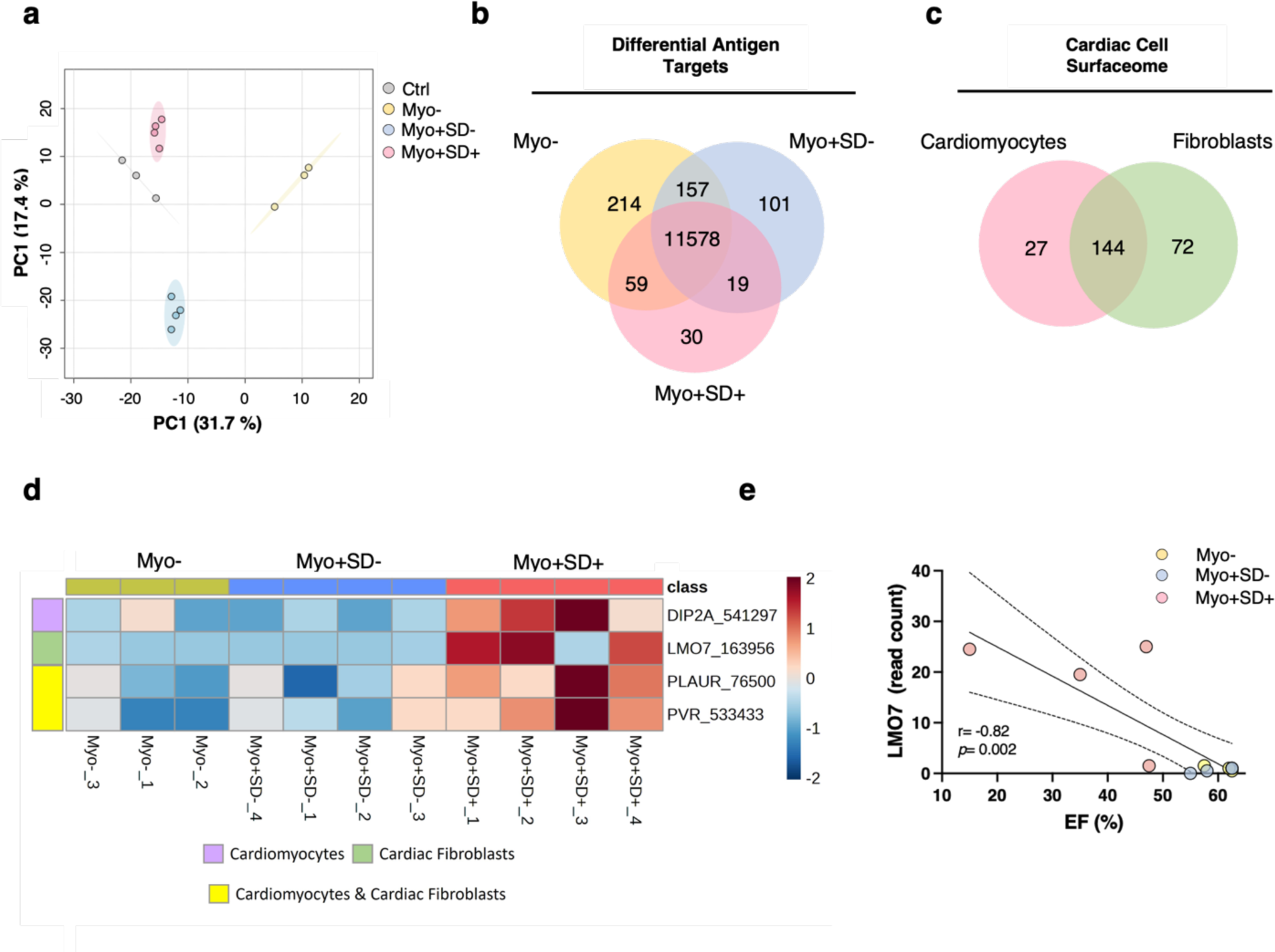
Identification of unique and potentially pathogenic autoantibody populations in Myo+SD+ patient serum. (**a**) PCA plot of differential antigen targets. (**b**) Venn diagram describing the number of antigen targets in each patient subgroup (**c**) Venn diagrams describing the number of cardiomyocyte and cardiac fibroblast surface proteins and their overlap. (**d**) Heatmap of potential pathogenic autoantibodies targeting cardiac cell surface antigens. (**e**) Linear correlation between patient EF and corresponding read counts of LMO7.

### Identification of candidate pathogenic autoantibodies

Based on the enhanced Myo+SD+ IgG binding to the cell surface, we sought to identify those specific targets. First, we characterized the cell surface proteome (surfaceome) of iPSC-CMs and primary cardiac fibroblasts by surface biotinylation, immunoprecipitation, and LC-MS/MS analysis. After data preprocessing and quality control filtering to omit intracellular proteins (Methods), we identified surface proteins unique to cardiomyocytes (n=27), cardiac fibroblasts (n=72) and shared by both cell types (n=143) (**Fig. 5c, Extended Data Fig. 10b, Supplementary Information Table 4**). By overlapping surface proteins with Myo+SD+ IgG-targeted antigens, we identified four candidate pathogenic autoantibodies. These autoantibodies targeted DIP2A, a cardiomyocyte-specific receptor associated with cardioprotection,^40^ LM07, a cardiac fibroblast-specific multifunctional regulator of protein-protein interactions,^41,42^ and 2 antigens on both cell types: PVR, associated with cell adhesion^40^, and PLAUR^43^, associated with extracellular matrix (ECM) degradation, **Fig. 5d, Supplementary Information Table 5**). Linear correlation analyses between these autoantibodies and EF indicated that autoantibodies to LMO7 had the strongest correlation (r= −.89, p=0.0002, **Fig. 5e**) with EF.

## DISCUSSION

We report a human cardiac tissue model of SLE and its use to study autoantibody-mediated heart injury. Using iPSC-derived cardiomyocytes, engineered human cardiac tissues, and patient-specific autoantibody repertoires, we identified two distinct signatures of autoantibody binding patterns that correlated with clinical phenotypes. *First*, IgG fractions from SLE patients with high levels of myocardial inflammation demonstrated increased binding to stressed cells within tissues subjected to electromechanical stimulation. *Second*, IgG from SLE patients with myocardial inflammation *and* systolic dysfunction (Myo+SD+) exhibited enhanced binding to the surface of viable cardiomyocytes. We showed that by binding to viable cardiac cells, IgG exerted potent biological effects independently of complement and immune cell involvement. These findings indicate that SLE autoantibodies evoke a range of biological processes that may contribute to clinical heterogeneity in disease manifestations and outcomes.

These experiments expanded upon our previously reported engineered cardiac tissue model^28^ by using metabolic maturation media in combination with an electromechanical stimulation regimen. Using isogenic hiPSCs for tissues enabled robustness and scalability but did not account for the diverse genetic backgrounds of SLE patients, an aspect worthy of further studies. Additionally, true autoreactivity was not demonstrated in this allogeneic model. However, the lack of binding of IgG from healthy controls to the engineered cardiac tissues suggests that the contribution of alloantibodies was minimal.

The intensity training regimen resulted in some apoptosis, potentially rendering the stressed tissue immunologically distinct due to the redistribution of intracellular antigens to the cell surface and allowing autoantibody penetration into apoptotic cells and their binding to intracellular entities. Consistent with previously reported autoantibody binding patterns in the heart, we found increased binding to the surface of apoptotic blebs and to the nucleus, highlighting that our model recapitulated the stressed cellular microenvironment reported in the myocardium of some SLE patients with myocardial inflammation^44,45^. The correlation between enhanced autoantibody binding to stressed cardiac cells and clinically elevated levels of myocardial inflammation suggests that autoantibody binding to stressed cardiac tissue may evoke immune cell infiltration and activation that leads to increased myocardial inflammation.

The lack of correlation between myocardial inflammation and systolic dysfunction suggests that autoantibody binding may directly influence cardiac tissue function beyond contributing to immune cell mediated inflammation. Building upon this observation and prior studies reporting that autoantibodies directly influences cell function despite the absence of immune cells^22,25^ we hypothesized that autoantibodies from patients with systolic dysfunction bind to the surface of viable cardiac cells and initiate downstream signaling or modulate physiological processes. The collected experimental evidence in this study supports this hypothesis. IgG purified from Myo+SD+ patients demonstrated enhanced binding to viable cardiomyocytes, leading to potent biological effects, including altered cellular composition, cardiomyocyte respiration, tissue calcium handling, and transcriptomics.

Transcriptomic analysis of cardiac tissues indicated downregulation of calcium channels, supporting the observation of impaired calcium handling in Myo+SD+ IgG-treated tissues. Upregulation of genes associated with mitochondria and ROS production was further confirmed by cellular assays. Interestingly, when cardiomyocytes were cultured with Myo+SD+ IgG there was a significant decrease in respiration and mitochondrial quality accompanied by an increase in mitochondrial quantity. This opposing directionality suggests that the effect of Myo+SD+ IgG to mitochondrial dysfunction is compensated, at least in part, by increased mitochondrial biogenesis. Interestingly, while most mitochondria associated genes were upregulated, two genes, *NDUFS4 and NDUFA12,* related to oxidative phosphorylation were downregulated in cardiomyocytes by the Myo+SD+ IgG, both of which are components of mitochondrial complex I. Mutational and knockout studies of *NDUFS4* have revealed mitochondrial dysfunction with compensatory increases in mitochondrial content as well as clinical and preclinical evidence of hypertrophic cardiomyopathy^46,47^, suggesting that impaired function of mitochondrial complex I might contribute both to our *in vitro* findings and to clinical observations. Transcriptomic data are also consistent with previous studies linking SLE autoantibodies to impaired cellular respiration and ROS production in other organ systems^48,49^. Moreover, impaired calcium handling, cytosolic calcium overload, and oxidative stress in the myocardium have all been linked to dysregulation of mitochondrial function, increased mitochondrial biogenesis, and the development of clinical systolic dysfunction^50–52^. Considering the abundant mitochondrial content in cardiomyocytes and the heart’s high metabolic demand, the perturbation in respiration and oxidative stress potentially renders the heart more vulnerable to IgG-induced injury compared to other organs and potentially leads to the development of systolic dysfunction.

The enhanced proliferation of cardiac fibroblasts incubated with Myo+SD+ IgGs suggests that this cellular response could contribute to the (*i*) late myocardial fibrosis and systolic dysfunction among neonatal patients with SLE-mediated congenital heart block^53^, and/or (*ii*) systolic dysfunction in SLE patients. As the specific phenotypes in cardiomyocytes and cardiac fibroblasts induced by Myo+SD+ IgG do not fully explain these and other clinical SLE phenotypes the study of autoantibodies on other cardiac cell populations (endothelial and smooth muscle cells, pericytes, and macrophages) may uncover valuable insights into disease pathogenesis.

The PhIP-seq analyses identified unique autoantibody profiles in SLE patients, including autoantibodies that were not previously reported in SLE. As anticipated, we did not discover a single autoantibody associated with myocardial injury. Instead, despite the wide array of autoantibodies in SLE, our studies revealed shared autoantibodies that were specific to Myo+SD+ patients. These findings indicate that a unique autoantibody repertoire in the Myo+SD+ patient subgroup contributes to the observed differential responses.

We identified potential pathogenic autoantibodies targeting the surface of cardiomyocytes and cardiac fibroblasts by combining PhIP-seq and cell surfaceome analyses. The cardiomyocyte-enriched IgG target, DIP2A, is a cell surface receptor for follistatin-like 1 (FSTL1), a protein that increases cardiomyocyte survival and decreases cardiomyocyte apoptosis and hypertrophy^40,54–58^. Thus, an autoantibody to the FSTL1 receptor may interfere with its cardioprotective role, and potentially contribute to the systolic dysfunction in the Myo+SD+ patients. The cardiac fibroblast specific protein LMO7 participates in the regulation of TGF-*β* and fibrotic remodeling^59^. Autoantibody-mediated inhibition of LMO7 could increase fibroblast activation and proliferation, accounting for our findings in IgG treated tissues. PLAUR, a urokinase plasminogen activator surface receptor has established roles in ECM proteolysis, a key process in fibrotic remodeling^60^, further connecting specific autoantibody targeting with fibrosis. Knockdown of PLAUR also impairs mitochondrial function, promotes immature biogenesis of mitochondria, and alters cell metabolism^61–63^, possibly contributing to the observed metabolic dysfunction. Further investigation of the mechanisms by which these candidate pathogenic autoantibodies contribute to systolic dysfunction and other antigen-antibody interactions may aid in the identification of new therapeutic targets to reduce SLE-associated myocardial injury.

Importantly, it has been shown that numerous intracellular autoantibodies cross-react with surface proteins on cardiomyocyte and alter their function. Therefore, the identified Myo+SD+ autoantibodies that target intracellular proteins might also play a role in the observed phenotype. In addition, studies of larger patient cohorts would advance autoantibody-phenotype associations, insights into molecular consequences, and allow combinatorial approaches for evaluating multiple candidate autoantibodies and distill individual and interactive effects on cardiac tissues.

In conclusion, our study demonstrates the utility of an engineered model of the human myocardium for elucidating the heterogeneity and pathophysiology of autoantibody mediated myocardial injury in SLE patients. We envision this model will improve myocardial risk stratification in SLE, facilitate the identification of new therapeutic targets and be extended to other diseases associated with complex autoantibody mediated tissue damage.

## DATA AVAILABILITY

All data are available in the main manuscript, extended data, or associated Figshare site. The original datasets are available from the corresponding author upon reasonable request.

## CODE AVAILABILITY

All codes utilized for the collection and/or analysis of data during the current study are available from the corresponding author upon reasonable request.

## ACKNOWLEDGMENTS

We gratefully acknowledge funding of this research by the National Institutes of Health (P41EB027062 and 3R01HL076485 to G.V-N.), the American Heart Association (19TPA34910217 to R.W.), a Pfizer Aspire research award (WI237809 2018 ASPIRE US Rheumatology to R.W.), and the National Science Foundation (NSF1647837 to G.V-N.). These studies used the Shared Resources of the Herbert Irving Comprehensive Cancer Center at Columbia University (HICCC), funded in part through the NIH/NCI Cancer Center Support Grant P30CA013696. We thank Michael Kissner and the staff of the Columbia Stem Cell Initiative Flow Cytometry Core Facility at Columbia University Irving Medical Center, Theresa Swayne and the staff of the Confocal and Specialized Microscopy Shared Resource at HICCC, and the staff of the Columbia Genome Center for their technical contributions to the work presented in this manuscript. We gratefully acknowledge the patients of the Lupus Center at Columbia University Irving Medical Center who contributed their serum samples and clinical data to this study. The authors declare no competing interests.

## EXTENDED DATA

**Extended Data Figure 1:**
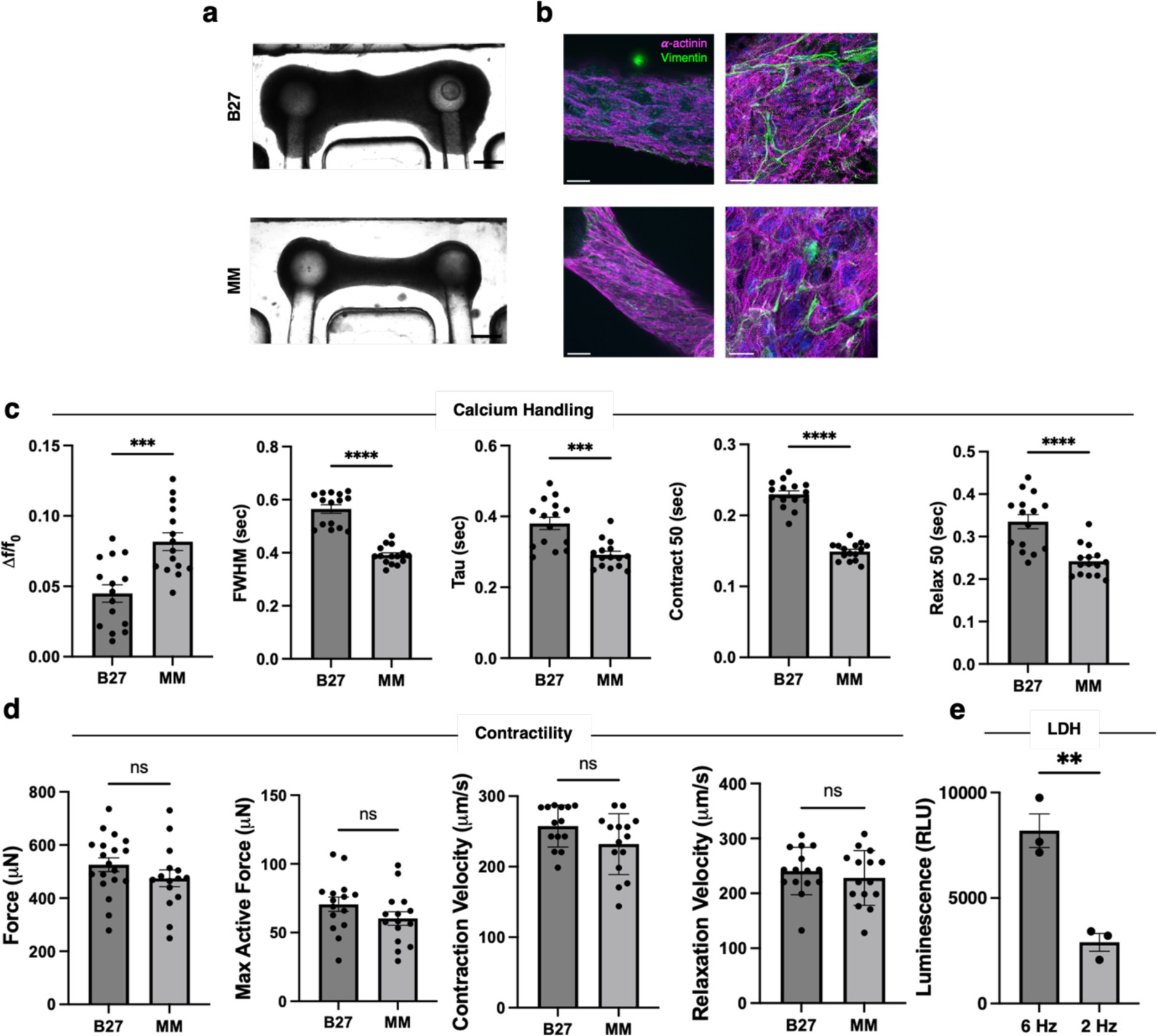
Characterization of engineered cardiac tissues cultured in different media formulations. (**a, b**) Representative brightfield (a) and immunofluorescence (b) images of engineered cardiac tissues cultured in standard RPMI-B27 (B27) medium (top) and metabolic maturation (MM) medium (bottom). (**c, d**) Measurements of calcium handling (c) and contractility (d) of engineered cardiac tissues cultured in B27 medium and MM. (**e**) Quantification of LDH release into supernatant from engineered cardiac tissues during electrical stimulation at 2 Hz and 6 Hz. Data are the mean ± s.e.m. Individual values are indicated by dots. B27, RPMI-1640 with B27 supplement. MM, metabolic maturation medium. LDH, lactate dehydrogenase. Scale bars in (a) are 500 μm; scale bars in (b) are 200 μm (left) and 10 μm (right).

**Extended Data Figure 2:**
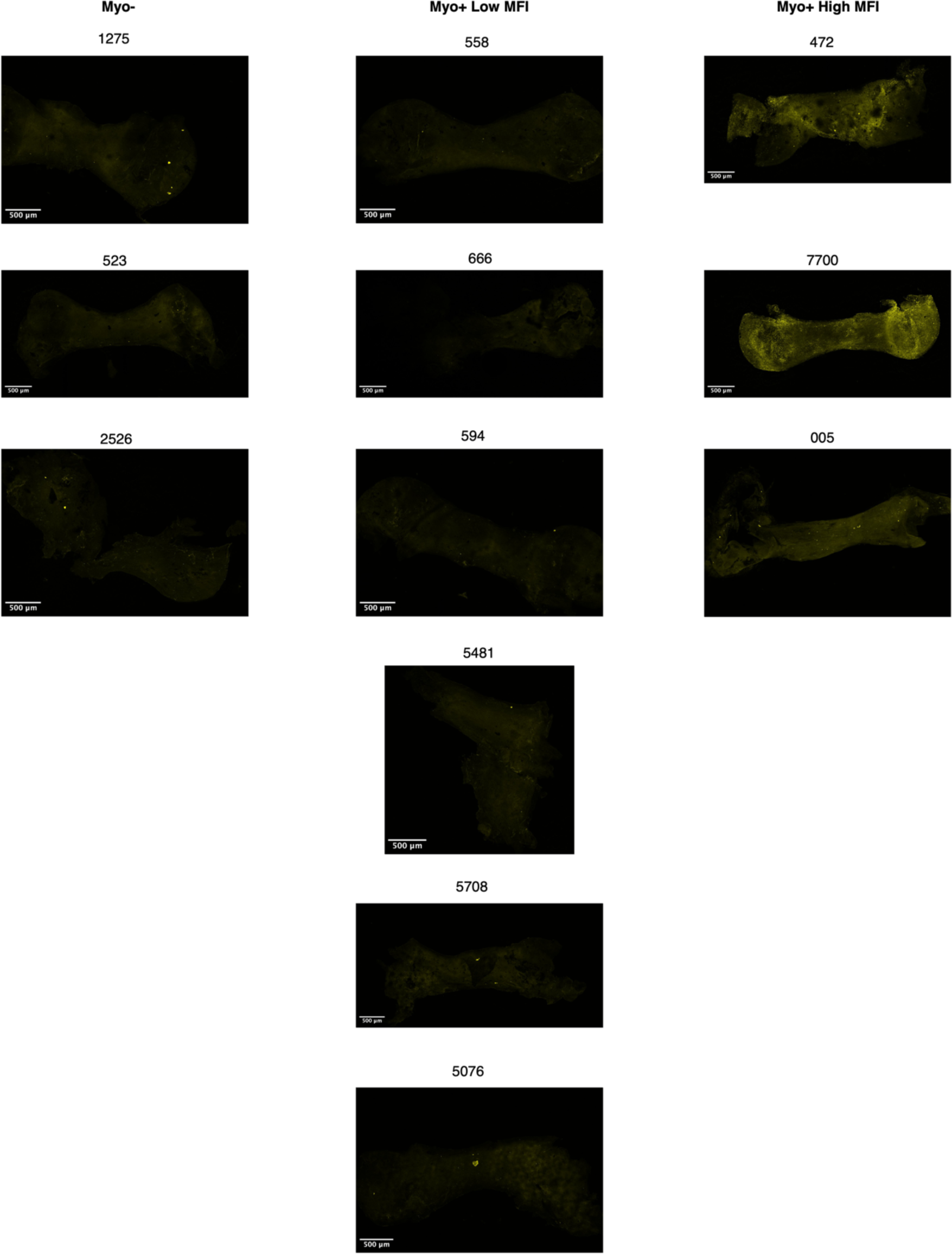
Indirect immunostaining of engineered cardiac tissues following treatment with patient IgG. Immunofluorescence images of engineered cardiac tissues treated with IgG isolated from each patient in the Myo-(left) and Myo+ (middle and right) groups, followed by staining with anti-human IgG to visualize patient-specific IgG binding (yellow). Patient samples from the Myo+ group are subclassified as low MFI (middle) and high MFI (right) based on quantification of human IgG staining.

**Extended Data Figure 3:**
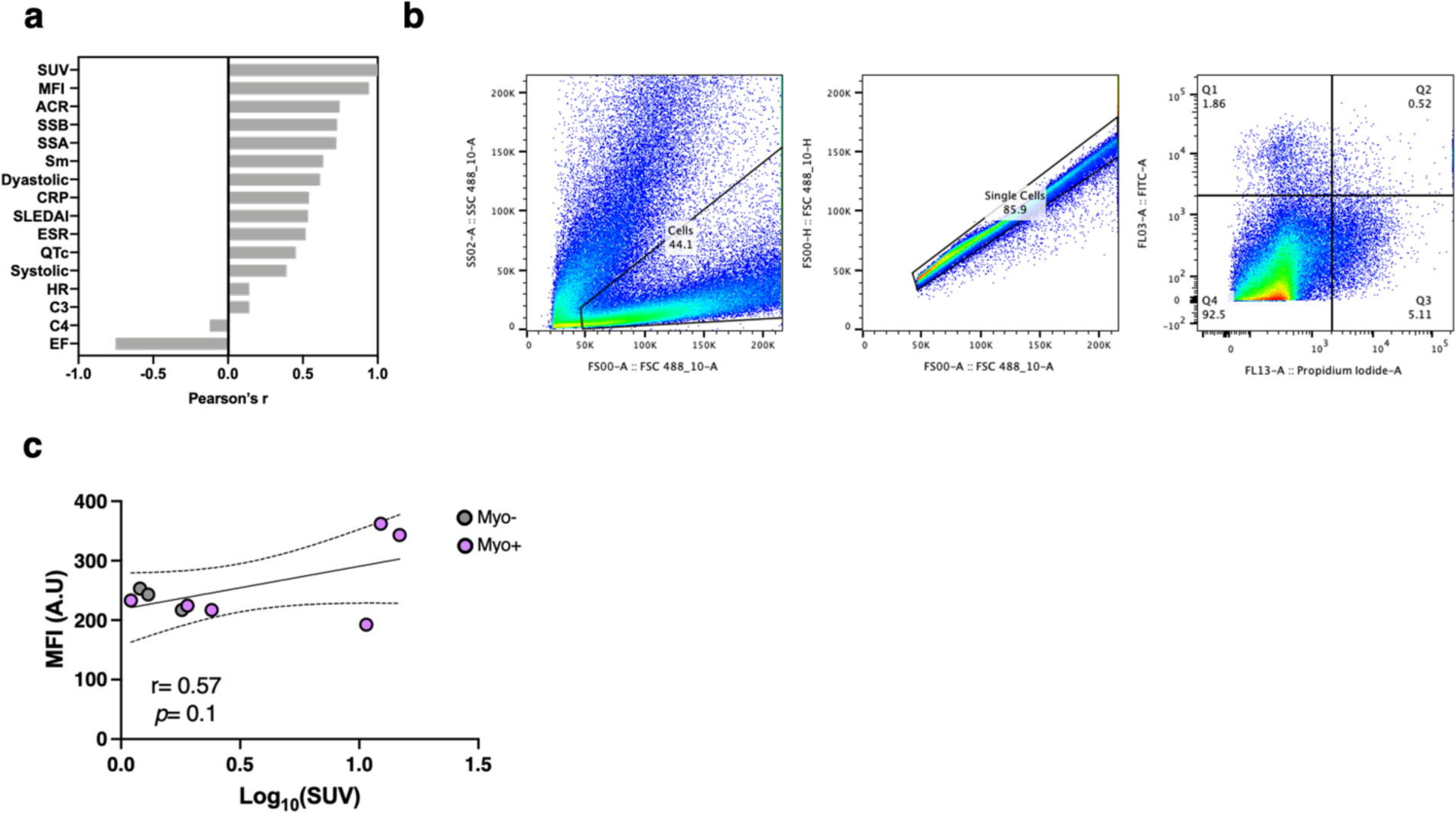
Quantitative analysis of IgG binding to engineered cardiac tissues and individual cells. (**a**) Summary of Pearson correlation coefficients of clinical ^18^F-FDG uptake quantified as SUVs compared with corresponding patient IgG binding levels to engineered tissues *in vitro* (MFI, second row) and with other clinical data collected as part of the cohort. (b) Example flow cytometry gating strategy used to identify viable cardiomyocytes, i.e., those that are double negative for propidium iodide and Apotracker Green (FITC). These are the cells in Q4. (**c**) Log-linear correlation between SUVs and MFI.

**Extended Data Figure 4:**
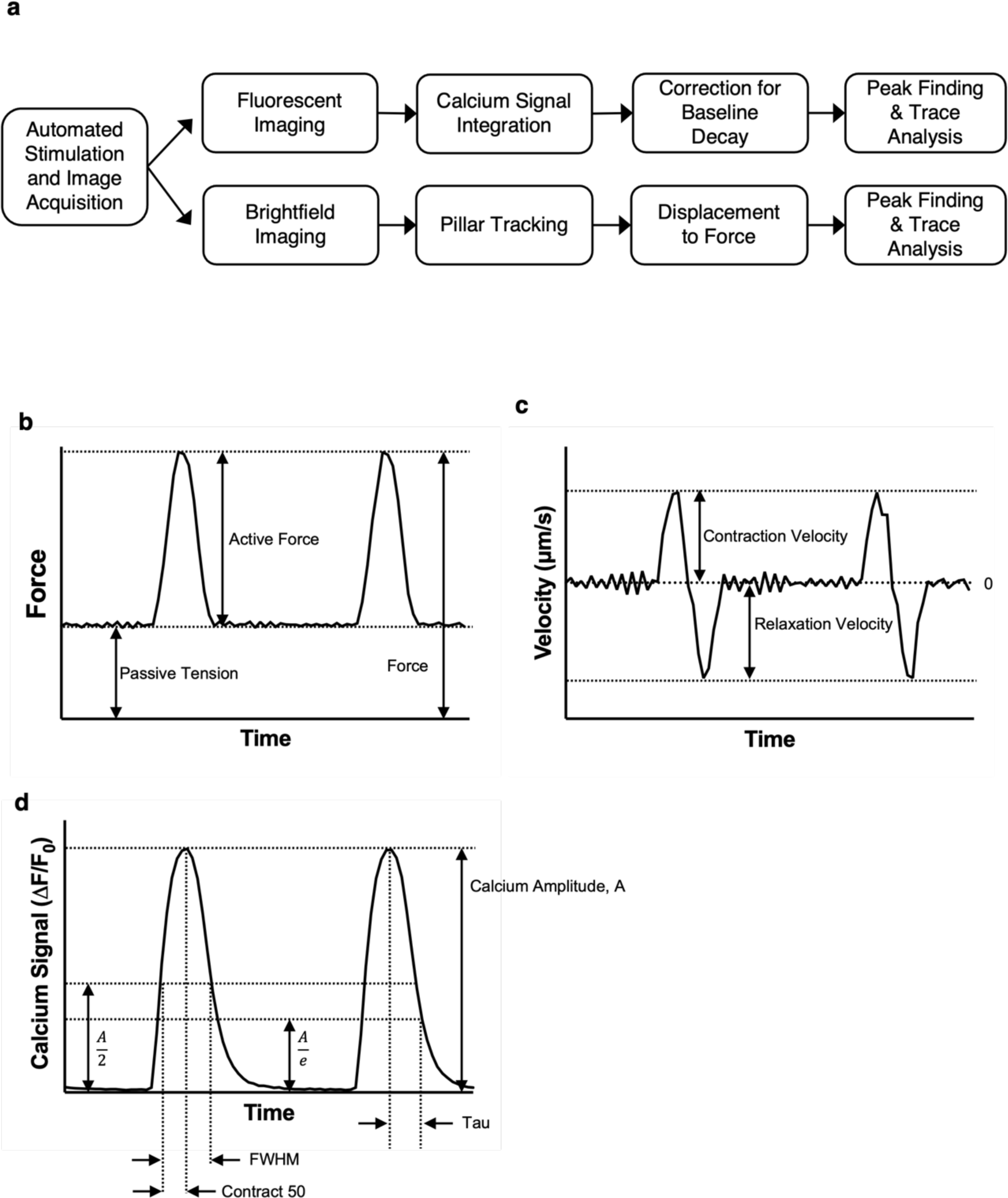
Functional analysis of engineered cardiac tissue. (**a**) Schematic overview of the automated imaging and analysis pipeline for the engineered cardiac tissues. (**b-d**) Representative traces of engineered cardiac tissue force generation (b), velocity (c), and calcium signal (d). Metrics used to quantify these traces are depicted on each plot.

**Extended Data Figure 5:**
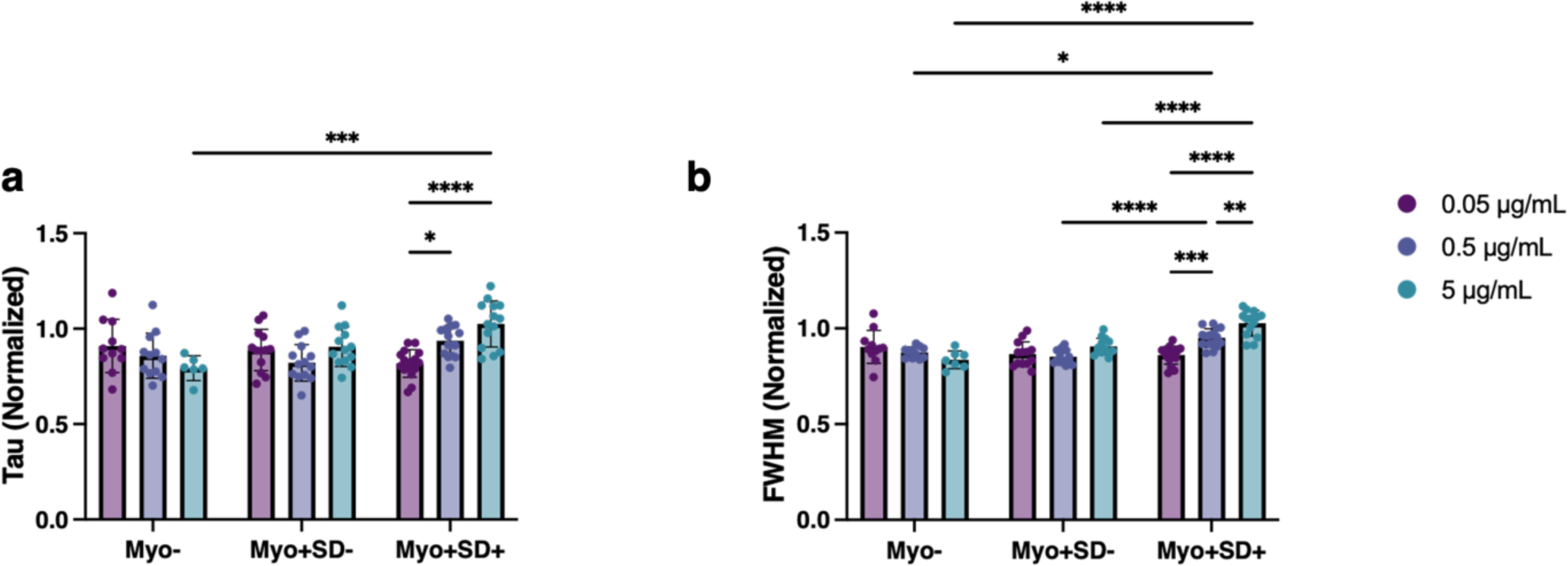
Dose-response effect of patient IgG on hiPSC-cardiomyocyte calcium handling. (**a,b**) IgG from Myo+SD+ patients demonstrate a clear dose-response effect in hiPSC-derived cardiomyocytes calcium handling measurements of tau (a) and FWHM (b). Data are the mean ± s.e.m. Individual values are indicated by dots; n = 18-24 independent hiPSC-cardiomyocyte samples per group (n = 3 technical replicates for each patient IgG sample); ordinary one-way ANOVA with Tukey’s test for multiple comparisons; * p < 0.05, ** p < 0.005, *** p < 0.001, **** p < 0.0001.

**Extended Data Figure 6:**
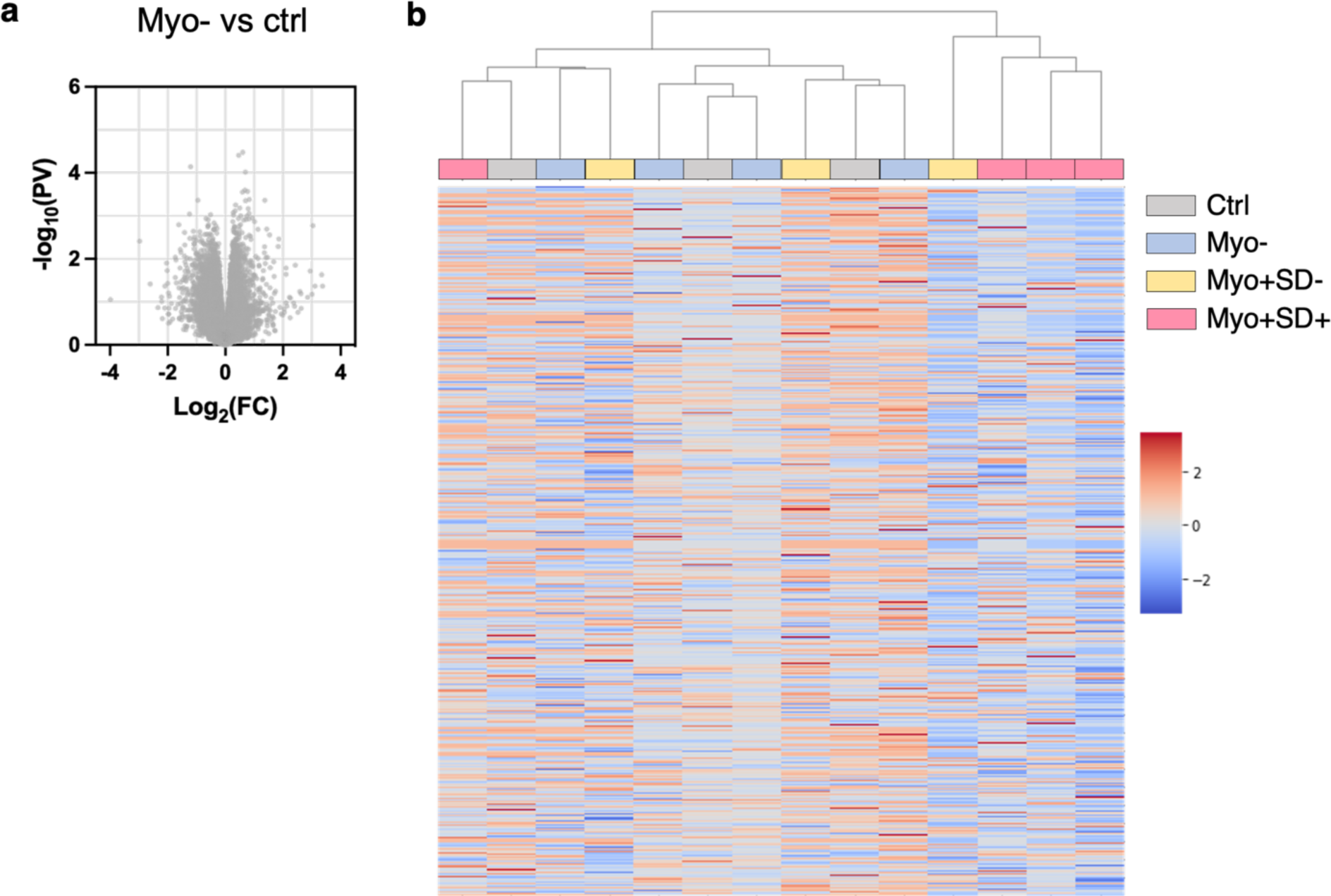
Transcriptomics of engineered cardiac tissues treated with patient IgG. (**a**) Volcano plot of DEGs between engineered cardiac tissues treated with SLE patient IgG and those treated with healthy control patient IgG. (**b**) Hierarchical clustering of overall transcriptomics of engineered cardiac tissues treated with IgG from each individual patient.

**Extended Data Figure 7:**
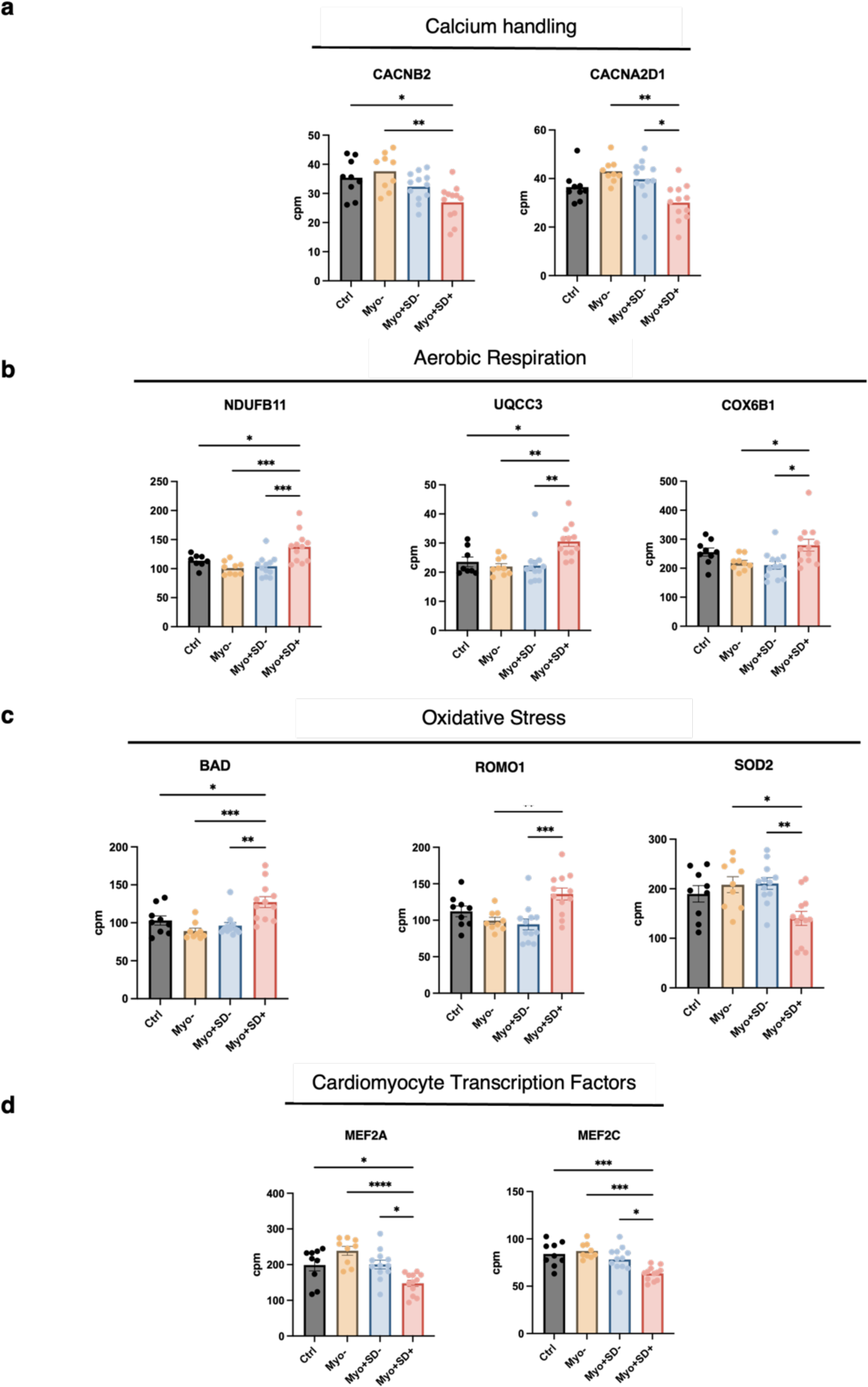
Gene expression measured by RNA-Seq for selected genes. Selected genes for calcium handling (a), Aerobic respiration (b), oxidative stress (c), and cardiomyocyte transcription factors (d). n = 3 technical replicates for each patient IgG sample; ordinary one-way ANOVA with Tukey’s test for multiple comparisons; * p < 0.05, ** p < 0.005, *** p < 0.001, **** p < 0.0001.

**Extended Data Figure 8:**
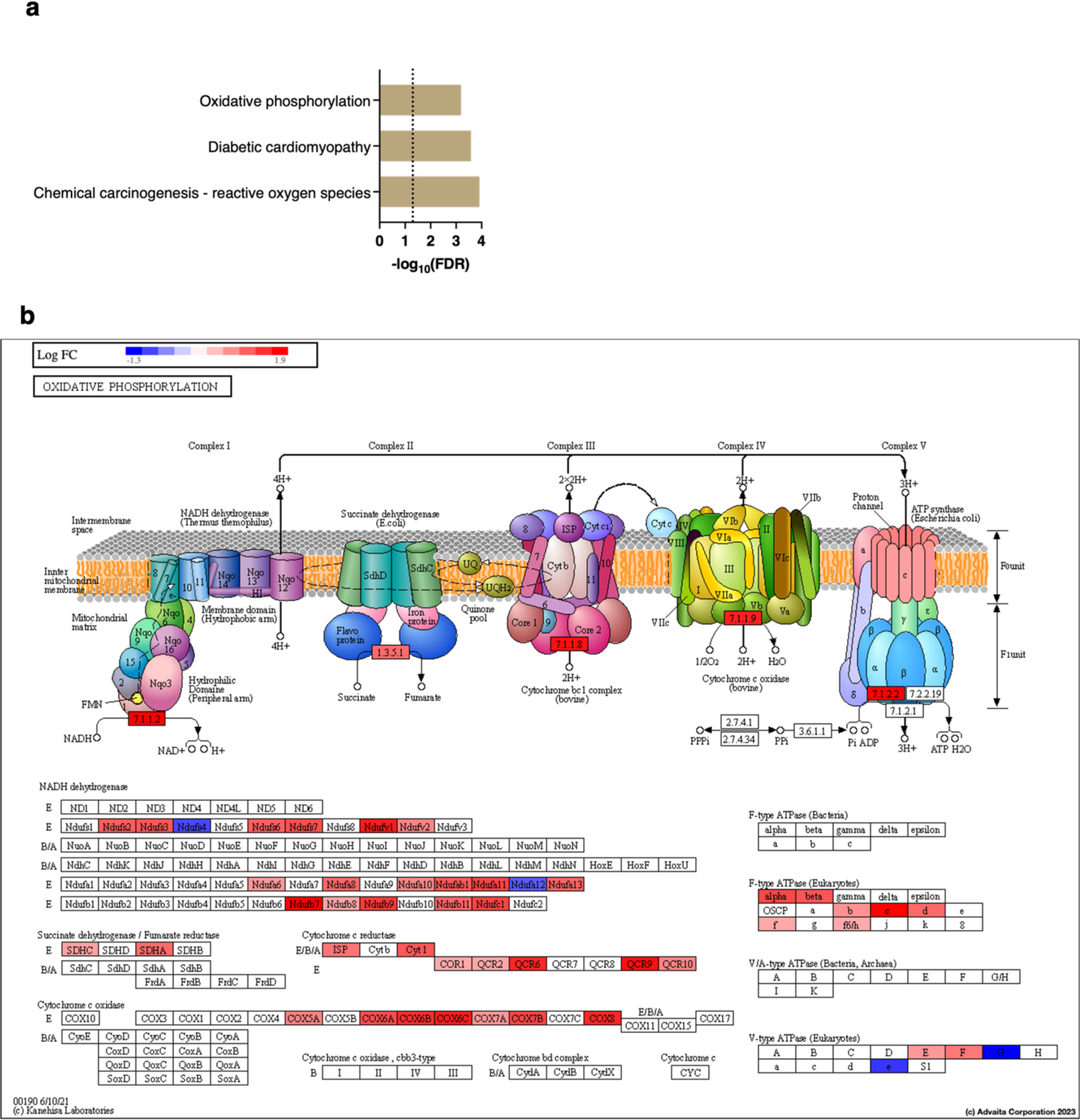
CIBERSORTx analysis of cardiomyocytes-specific gene expression profiles. (**a**) KEGG, pathway analysis of cardiomyocytes-specific DEGs between the Myo+SD+ and the Myo-subgroups. (**b**) Schematic of KEGG oxidative phosphorylation pathway. Colored boxes indicate differentially expressed genes between the Myo+SD+ and the Myo-subgroups.

**Extended Data Figure 9:**
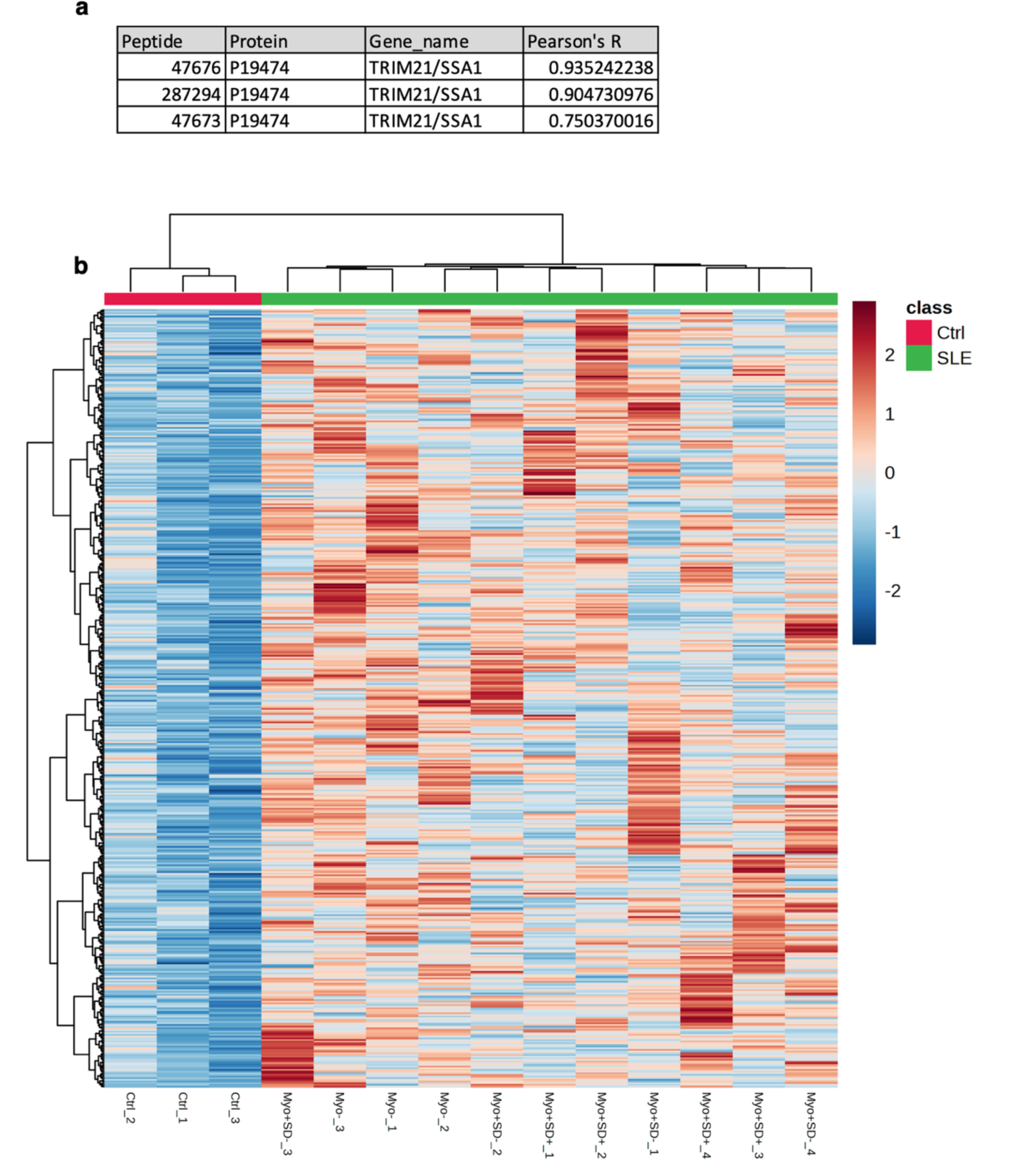
PhIP-Seq analysis of patient-specific antigen targets. (**a**) Table summarizing Pearson’s R quantified by correlation between different SSA peptides measured by PhIP-Seq and clinical laboratory measurements of patient SSA antibodies. (**b**) Hierarchical clustering of peptides increased in SLE compared to healthy controls. Each column represents a patient (green; SLE patients, red; healthy controls). Each row represents a peptide identified by PhIP-Seq.

**Extended Data Figure 10:**
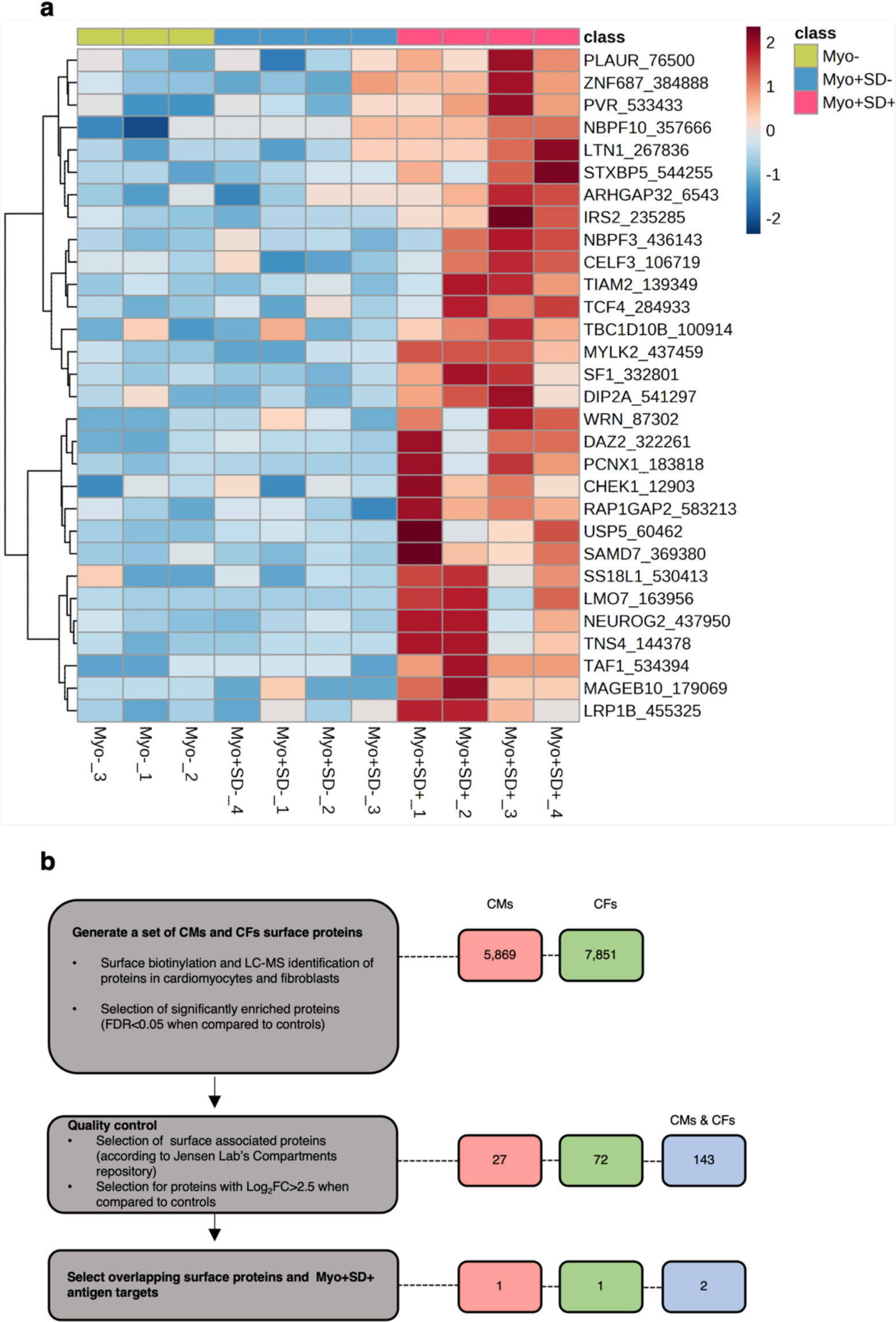
PhIP-Seq analysis of Myo+SD+ epitope targets and cell surface characterization. (**a**) Hierarchical clustering of peptides increased in Myo+SD+ patients compared to Myo- and Myo+SD-patients. Each column represents a patient (yellow: Myo-patients; green: Myo+SD-patients; purple: Myo+SD+ patients). Each row represents an identified epitope. 76 epitopes from 65 proteins. (**b**) Schematic of surface protein analysis pipeline and quantification of identified proteins.

## SUPPLEMENTARY INFORMATION

**Supplementary Information Table 1: Differentially expressed genes in engineered cardiac tissues treated with SLE patient IgG. (a)** Myo+SD+ vs Myo+SD-**(b)** Myo+SD+ vs Myo-

**Supplementary Information Table 2: CIBERSORTx analysis of cardiomyocyte-specific differentially expressed genes in engineered cardiac tissues treated with Myo+SD+ IgG. (a)** Myo+SD+ vs Myo+SD-(b) Myo+SD+ vs Myo-

**Supplementary Information Table 3: Phage Immunoprecipitation Sequencing. (a)** Peptides with increased IgG reactivity in serum of systemic lupus erythematosus (SLE) patients compared to control (CTRL). **(b)** Peptides with different IgG reactivity in serum of Myo+SD+ patients compared with Myo+SD-patients. **(c)** Peptides with different IgG reactivity in serum of Myo+SD+ patients compared to Myo-patients. **(d)** Peptides with different IgG reactivity in serum of Myo+SD-patients compared to Myo-patients. **(e)** Peptides with different IgG reactivity in serum of Myo+SD+ patients compared to patients without systolic dysfunction or without myocarditis (combined Myo+SD- and Myo-groups).

**Supplementary Information Table 4: Surface protein characterization.** Total identified surface proteins. Columns list the proteins identified on the surface of both hiPSC-cardiomyocytes and primary cardiac fibroblasts (iPS-CMs & CFs), only on hiPSC-cardiomyocytes (iPS-CMs), and only on primary cardiac fibroblasts (CFs), respectively.

**Supplementary Information Table 5: Potential disease contributing autoantibodies**. Expression levels of each cognate antigen are included from the Human Heart Cell Atlas^39^.

## METHODS

### Patient cohort

All study participants signed an informed consent form. The study was approved by the Columbia University Institutional Review Board. Patients were recruited from the Columbia University Lupus Cohort in 2 groups: one investigating the presence of myocardial inflammation (defined by FDG-PET myocardial uptake) in SLE patients without active cardiac symptoms, and the other investigating SLE patients evaluated with FDG-PET following active cardiac symptoms. Patients were 18 years of age or older and met the 1997 American College of Rheumatology classification criteria for SLE^31^. SLE patients with both myocardial inflammation and systolic dysfunction were only included in the study if review of previous EF measurements revealed an acute decline in EF at the time of enrollment and serum collection.

### Clinical covariates

SLE disease duration was defined as the duration in years from the date of physician diagnosis. SLE disease activity was calculated using the Systemic Lupus Erythematosus Disease Activity Index 2000 (SLEDAI-2K)^64^. Traditional cardiovascular risk factors, history of lupus nephritis and antiphospholipid antibody syndrome, as well as use of immunosuppressants was ascertained by patient questionnaire and medical record review.

### Laboratory covariates

Autoantibodies including antinuclear antibodies, anti-SSA/Ro, anti-SSB/La, anti-ds-DNA, anti-Smith, anti-RNP, antiphospholipid antibodies and other pertinent laboratories such as complement levels (C3, C4), erythrocyte sedimentation rate (ESR), high-sensitivity C reactive protein, troponin and pro-beta-natriuretic peptide (pro-BNP) levels, were assessed at the clinical laboratory at New York Presbyterian Hospital and the Core Laboratory of the Columbia University Irving Institute for Clinical and Translation Research.

### ^18^F-FDG PET/CT

Myocardial uptake imaging was performed on an MCT 64 PET/CT scanner (Siemens Medical Solutions USA, Knoxville, Tennessee, USA). A low-dose CT transmission scan (120 kV, 25 mA) was obtained for attenuation correction of PET data. All patients were on a carbohydrate-free diet for 24 hours. Patients were injected with 10±0.1 mCi of ^18^F-FDG intravenously using an antecubital or dorsal forearm catheter. A list mode 3D PET scan was acquired for 10 min following a 90 min uptake period post-^18^F-FDG injection. Non-gated attenuation-corrected images were reconstructed yielding 3 mm effective resolution. Corridor 4DM software was used to visually assess myocardial ^18^F-FDG uptake as well as semi-automatically quantify mean radiotracer uptake in the myocardium. Quantification of inflammation by ^18^F-FDG PET/CT involved measurement of SUV in the myocardium. SUV was determined for 17 myocardial segments and the maximum of these values was determined as SUVmax.

### Isolation of patient IgG fractions

Samples of serum were collected from patients at the time of ^18^F-FDG PET/CT scans and stored at −80°C until IgG purification. The IgG antibody fractions were isolated from patient sera by affinity purification with immobilized Protein A/G using the NAb Protein A/G Spin Kit (Thermo Fisher Scientific, cat. no. 89950), according to the manufacturer’s protocol. Following, Thermo Scientific™ Zeba™ spin desalting columns (Thermo Fisher Scientific, cat. no. 87766) were used to further purify IgG from contaminates. Importantly, IgG purification from the patients’ sera was performed to eliminate confounding effects of sera components such as cytokines, growth factors and drugs. Purified IgG fractions were stored at −80°C until experimental use.

### hiPSC sourcing

The experiments were performed using the WTC11-GCaMP6f hiPSC line, obtained through an institutional Materials Transfer Agreement (MTA) with the Gladstone Institutes (Dr. Bruce Conklin) ^65^. The line contains a constitutively expressed GCaMP6f calcium-responsive fluorescent protein ^66^ inserted into a single allele of the *AAVS1* safe harbor locus, which enables real-time and label-free visualization of calcium handling. The parental hiPSC line, WTC11 (GM25256 at the Coriell Institute), was generated from a healthy donor with no history of cardiovascular or autoimmune diseases ^67^.

### hiPSC culture

hiPSCs were cultured in mTeSR Plus medium (STEMCELL Technologies, cat. no. 100-0276) on tissue culture plates coated with Matrigel (Corning, cat. no. 354230; diluted 1:100) and passaged every three to four days with 0.5 mM EDTA (Thermo Fisher Scientific, cat. no. 15575) in PBS (Corning, cat. no. 21-040). For the first 24 hours after passaging, 5 μM Y-27632 dihydrochloride (Tocris Bioscience, cat. no. 1254), a rho-associated protein kinase (ROCK) inhibitor, was added to the culture medium. hiPSCs were karyotyped and regularly tested for mycoplasma contamination.

### Cardiomyocyte differentiation from hiPSCs

Cardiomyocytes were differentiated as previously described^68^. Briefly, hiPSCs were plated at a density of 155,000 cells/cm^2^ two days prior to the start of differentiation. Culture medium was replaced with cardiomyocyte differentiation medium (CDM), consisting of RPMI-1640 medium (Thermo Fisher Scientific, cat. no. A4192302), 500 μg/mL recombinant human albumin (Sigma-Aldrich, cat. no. A9731), and 213 μg/mL L-ascorbic acid 2-phosphate (Sigma-Aldrich, cat. no. A8960). From day 0 to day 2, CDM was supplemented with 4 μM CHIR99021 (Tocris Bioscience, cat. no. 4423), and from day 2 to day 4, CDM was supplemented with 2 μM Wnt-C59 (Tocris Bioscience, cat. no. 5148). Following CDM was changed every two days. On day 10, CDM was replaced with purification medium, consisting of RPM-1640 without glucose (Thermo Fisher Scientific, cat. no. 11879020), 1x B27 Supplement (Thermo Fisher Scientific, cat. no. 17504044), and 213 μg/mL L-ascorbic acid 2-phosphate, to purify the hiPSC-cardiomyocytes population and eliminate potential contaminating mesodermal and endodermal populations. On day 13, the medium was replaced with RPMI-B27 medium, consisting of RPMI-1640 medium, 1xB27 Supplement, and 213 μg/mL L-ascorbic acid 2-phosphate. On day 17, cells were pretreated with 5 μM Y-27632 dihydrochloride for 4 hours to prepare for dissociation. Cells were dissociated by enzyme digestion with 95 U/mL collagenase type II (Worthington, cat. no. LS004176) and 0.6 mg/mL pancreatin (Sigma–Aldrich, cat. no. P7545) in a dissociation buffer consisting of 5.5 mM glucose, 1.8 mM CaCl_2_, 5.4 mM KCl, 0.81 mM MgSO_4_, 100mM NaCl, 44 mM NaHCO_3_, and 0.9 mM NaH_2_PO_4_. Cells were incubated at 37° C on a shaker for approximately 30 minutes. Flow cytometry for cardiac troponin T (cTnT; BD BioSciences cat. no 565744) was performed prior to cell use for tissue fabrication to ensure cell purity (>90% cTnT+).

### Cardiac fibroblasts

Primary human ventricular cardiac fibroblasts (NHCF-V; Lonza, cat. no. CC-2904) were cultured according to the manufacturer’s protocol with Fibroblast Growth Medium 3 (PromoCell, cat. no. C-23130). Fibroblasts were used for engineering tissues between passage 3 and passage 5. Cells were generated from healthy donors with no history of SLE or cardiovascular disease.

### milliPillar Platform fabrication

The platform, termed milliPillar, was assembled as previously described^28^. Briefly, the platform was fabricated by casting polydimethylsiloxane (PDMS; Dow Chemical Company, cat. no. 02065622; 1:10 curing agent to elastomer ratio) into custom molds containing carbon electrodes and curing overnight at 65° C. The platform was then detached from the mold and plasma bonded to a glass slide. After completion, each well contained a set of horizontal flexible pillars upon which engineered tissues can be suspended, as described below. Each well also contained a set of carbon electrodes for electrical field stimulation. A custom Arduino-based electrical stimulator was used for electrical pacing during culture and video acquisition, as previously described^28^.

### Human engineered cardiac tissues

Engineered cardiac tissues were fabricated as previously described^28^. Briefly, dissociated hiPSC-Cardiomyocytes and primary cardiac fibroblasts were mixed in a 3:1 ratio and resuspended in fibrinogen by mixing the cell solution with 33 mg/mL human fibrinogen (Sigma-Aldrich, cat. no. F3879) and RPMI-B27 (described above) to a final fibrinogen concentration of 5 mg/mL and cell concentration of 37,000 cells/μL. A 3 μL drop of 2.5 U/mL human thrombin (Sigma-Aldrich, cat. no. T6884) was added to each well of the platform, followed by addition of 12 μL of the cell-fibrinogen solution. The two solutions were mixed, spread evenly across the well, and incubated 37°C for 15 min, allowing the fibrin hydrogel to polymerize within the well. Following polymerization, each well was filled with 400 µL of RPMI-B27 supplemented with 10µM Y-27632 and 5 mg/mL 6-aminocaproic acid (Sigma-Aldrich, cat. no. A7824). On day 1, RPMI-B27 medium was replaced without Y-27632.

### Optimization of engineered cardiac tissue culture medium

On day 2, platforms were randomized into two different culture medium groups: RPMI-B27 and a previously described metabolic maturation medium (“MM”)^33^. Medium was changed every other day. On day 6, medium was replaced without 6-aminocaproic acid. On day 7 electrical stimulation was initiated at a frequency of 2 Hz until day 14. Tissue functionality was assessed by calcium and brightfield imaging on day 14. Force generation, contraction and relaxation velocities, and calcium transient amplitude and kinetics were measured (**Extended Data Fig. 4**). Tissues cultured in MM displayed enhanced compaction (**Extended Data Fig. 1b**), cell alignment, α-actinin striations (**Extended Data Fig. 1c**), and calcium handling (**Extended Data Fig. 1d**). The contractile behavior was comparable between tissues cultured in MM and RPMI-B27 (**Extended Data Fig. 1e**). MM was chosen as the culture medium for subsequent experiments with patient IgG.

### Engineered cardiac tissue incubation with autoantibodies

Patient-specific serum purified IgG was added to the culture medium on day 14 of tissue cultivation, with tissues stimulated at 2 Hz and maintained for an additional 14 days. During this time, the frequency of electrical stimulation was stepwise increased from 2 to 6 Hz by day 21 followed by decrease to 2 Hz until day 28 (intensity training). To exclude the possibility that the functional effect on cardiac tissue performance is not associated to titer levels and only to distinct autoantibody patterns, the same concentration of purified patient autoantibodies, 0.5 μg/mL was added to all tissues. To assess the effect of purified IgG on cardiac tissue functionality we evaluated the tissue contractility and calcium handling (the full list of metrics and their descriptions is provided in **Extended Data Fig. 4**) at baseline (day 14), and after the addition of IgG, on days 21 and 28. To reduce noise from tissue-to-tissue variability, measurements were normalized to the baseline.

### Immunostaining

Whole mount engineered cardiac tissues were fixed and permeabilized in 100% ice cold methanol for 10 min, washed three times in PBS, and then blocked for 1 hour at room temperature in PBS with 2% fetal bovine serum (FBS). After blocking, the tissues were incubated with primary mouse anti–α-actinin (sarcomeric) antibody (1:750, Sigma-Aldrich, cat. no. A7811), and vimentin (Abcam, cat. no. ab24525) washed three times and incubated for 1 hr with secondary antibodies (Millipore Sigma, cat. no. AP194C; Thermo Fischer, cat. no. A-21206; Thermo Fischer, cat. no. A-31571). For nuclei detection, the tissues were washed and incubated with NucBlue (Thermo Fisher, cat. no. R37606). Whole tissues were placed in incubation chambers (Grace Bio-Labs, cat. no. 645501) and mounted with ProLong Diamond antifade mountant (Invitrogen, cat. no. 36961). Samples were visualized using a scanning laser confocal microscope (A1 confocal system with Eclipse Ti inverted microscope, Nikon Instruments).

### Indirect immunostaining

Engineered cardiac tissues and hiPSC-cardiomyocytes cultured with patient IgG were fixed and incubated with secondary anti-human IgG (Thermo Fisher Scientific, cat. no. A-11013) to visualize cellular autoantibody targets. Importantly, prior to fixation the tissues/cells were either permeabilized to detect all extracellular and intracellular binding sites or not permeabilized to detect IgG binding sites on the cell surface. Samples were visualized using a scanning laser confocal microscope (A1 confocal system with Eclipse Ti inverted microscope, Nikon Instruments). Binding intensity was measured by image analysis in ImageJ.

### Flow cytometry to assess autoantibody binding

iPSC-derived cardiomyocytes were differentiated and dissociated as described above. Individual patient IgG samples were diluted 1:500 in cell staining buffer (BioLegend, cat. no. 420201) and incubated with cardiomyocytes for 1 hour at room temperature, followed by incubation with CF-647 conjugated anti-human IgG (Sigma Aldrich, cat. no. SAB4600181) diluted 1:500 in cell staining buffer for 30 minutes at room temperature. Cardiomyocytes were then stained with Apotracker Green (BioLegend, cat. no. 427401) and propidium iodide (ThermoFisher, cat. no. R37169) according to the manufacturers’ protocols. Flow cytometry was conducted with a ZE5 Cell Analyzer (BioRad) to identify viable cardiomyocytes and quantify IgG binding. FlowJo software (BD Biosciences) was used for analysis. Viable cardiomyocytes were defined as those with negative staining for propidium iodide, a marker of dead cells, and Apotracker Green, a marker of apoptotic cells. An example of the gating strategy is provided in **Extended Data Fig. 3b**. IgG binding was quantified as mean fluorescent intensity for all cells within the viable cardiomyocyte population.

### Calcium imaging and analysis

To decrease tissue-to-tissue variability, all metrics were measured as percent change from baseline. Tissues were imaged in a live-cell chamber (STX Temp & CO_2_ Stage Top Incubator, Tokai Hit) using a sCMOS camera (Zyla 4.2, Andor Technology) connected to an inverted fluorescence microscope with a standard GFP filter set (IX-81, Olympus). Tissues were then electrically stimulated at 1 Hz, and videos were acquired at 100 frames per second (fps) for 300 frames to measure calcium flux. Calcium signals were analyzed as previously described^28^. Briefly, a custom Python script was developed to average the pixel intensities for each frame, and this transient was then corrected for fluorescent decay. Further analysis by the script then extracted the metrics of calcium handling including calcium amplitude, full width half max (FWHM), contract 50, and the exponential decay constant (*τ*). A description of these metrics is provided in **Extended Data Fig. 4**.

### Contractile function

For force generation measurements, videos were acquired at 20 fps for 4800 frames using a custom program to stimulate cardiac tissues from 0.5 Hz to 4 Hz as previously described, and force generation was analyzed from brightfield videos as previously described^28^. Briefly, a custom Python script was developed to track the motion of the pillar heads and to calculate the force by multiplying the displacement of the pillars with the coefficient determined from the force-displacement calibration curve generated for the pillars. The script used the location of the pillar heads to determine the total deflection of the pillar from their equilibrium position without any force applied. Further analysis of the deflection trace extracted metrics of contractility, including the following: force, passive force, active force, contraction velocity, and relaxation velocity. A description of these metrics is provided in **Extended Data Fig. 4**. To decrease tissue-to-tissue variability, all metrics were measured as percent change from baseline.

### Analysis of tissue function

Following feature extraction, tissues that did not capture at 1 Hz stimulation based on both calcium and contractility analysis at baseline were removed from further analysis. The data was then processed to remove outliers greater than or less than +/- 1.5 times the interquartile range for each group. Outliers were removed from each metric independently. The metrics for each tissue were internally normalized to baseline values and presented as percent change from baseline. For correlation analyses, Pearson’s correlation coefficients were calculated and reported.

### Surface protein analysis

Human primary cardiac fibroblasts were seeded in 10 cm dishes and hiPSC-Cardiomyocytes were plated in 6-well plates. Two wells were used per pulldown reaction. The Pierce Cell Surface Biotinylation and Isolation Kit (Thermo Fisher Scientific, cat. no. A44390) was used per the manufacturer’s protocol to biotinylate and pulldown cell surface proteins with minor alterations. Prior to biotinylation, cells were washed 4 times with PBS. Biotinylation reaction was conducted for 30 minutes at room temperature after which the cells were washed 4 times with PBS and lysed on the plate in the provided lysis buffer with 1x Halt Protease and Phosphatase Inhibitor Cocktail (Thermo Fisher Scientific, cat. no. 78442). Protein concentrations were determined with BCA assay. An equivalent amount of 1000 ug protein was used per pulldown reaction, and cell lysate was incubated with streptavidin beads for two hours at room temperature. The surface proteins were eluted in 40 μL custom elution buffer (12.5 mM biotin, 7.5 mM HEPES (pH 7.5), 75 mM NaCl, 1.5 mM EDTA, 0.15% (w/v) SDS, 0.075% (w/v) Sarkosyl, and 0.02% (w/v) Na-Deoxycholate) at room temperature for 30 minutes and an addition 40 μL custom elution buffer at 65°C for 20 minutes. Subsequently, 30 μL was used for Western blot confirmation and the remaining 50 μL was analyzed with mass spectrometry.

### Global surface protein quantitative analysis

For global surface protein quantitative analysis of cardiac fibroblasts and cardiomyocytes, diaPASEF-based method was used^69^. In brief, the eluted proteins from streptavidin beads were denatured in SDC buffer (1% SDC, 100 mM TrisHCl pH 8.5) and boiled for 20 min at 60°C, 1000 rpm. Protein reduction and alkylation of cysteines were performed with 10 mM TCEP and 40 mM CAA at 45° C for 20 min followed by sonication in a water bath, cooled down to room temperature. Proteins were precipitated with the SP3 method, as previously described^70^, and SP3 beads were resuspended in SDC buffer (1% SDC /100mM Tris-pH 8.5). Protein digestion was processed overnight by adding LysC / trypsin mix in a 1:50 ratio (µg of enzyme to µg of protein) at 37°C and 1400 rpm. Peptides were acidified by adding 1% TFA, vortexed, subjected to StageTip clean-up via SDB-RPS2, and dried in a speed-vac. Peptides were resuspended in 10 μL of LC buffer (3% ACN/0.1% FA). Peptide concentrations were determined using NanoDrop and 200 ng of each sample were used for diaPASEF analysis on timsTOFPro.

### LC-MS/MS for surface protein characterization

Peptides were separated within 87 min at a flow rate of 400 nl/min on a reversed-phase C18 column with an integrated CaptiveSpray Emitter (25 cm x 75 μm, 1.6 μm, IonOpticks). Mobile phases A and B were with 0.1% formic acid in water and 0.1% formic acid in ACN. The fraction of B was linearly increased from 2 to 23% within 70 min, followed by an increase to 35% within 10 min and a further increase to 80% before re-equilibration. The timsTOF Pro was operated in diaPASEF mode and data was acquired at defined 22 × 50 Th isolation windows from m/z 440 to 1,440. Two IM windows per diaPASEF scan were set. The collision energy was ramped linearly as a function of the mobility from 59 eV at 1/K0=1.6 Vs cm-2 to 20 eV at 1/K0=0.6 Vs cm-2. The acquired diaPASEF raw files were searched with the UniProt Human proteome database (UP000005640) in the DIA-NN search engine with default settings of the library-free search algorithm^71^. DIA-NN performed the search trypsin digestion with up to 2 missed cleavages. The following modifications were used for protein identification and quantification: carbamidomethylation of cysteine residues (+57.021 Da) was set as static modifications, while acetylation of the protein N-terminus (+42.010), oxidation of methionine residues (+15.995 Da) was set as a variable modification. The false discovery rate (FDR) was set to 1% at the peptide precursor and protein level. Results obtained from DIA-NN were further analyzed using Perseus software^72^. Significantly changed protein candidates were determined by t-test with a threshold for significance of p < 0.05 (permutation-based FDR correction) and 0.58 log2FC. Since isolation procedures may yield some contaminating cells with compromised plasma membranes, further validation step of the significant proteins was performed and data was compared to three subcellular localization data sources; Human Protein Atlas^73^, Jensen Lab Compartment Repository^74^, and CellSurfer^75^ for further quality control (**Extended Data Figure 7a**).

### RNA isolation

Tissues were snap frozen in liquid nitrogen and stored at −80° C until use. To isolate RNA, snap-frozen tissues were added to 300 μL of RNA lysis buffer with a stainless-steel bead (BioSpec, cat. no. 11079123) and homogenized with a Mini-Beadbeater (Biospec, cat. no. C321001) for 5-10 seconds or until tissue samples were not visible. RNA was then isolated using a Qiagen RNeasy Micro Kit (Qiagen, cat. no. 74004), according to manufacturer’s instructions. RNA for each sample was then analyzed for quality/quantity using an Agilent bioanalyzer, Qubit 2.0 at Columbia’s Molecular Pathology core.

### RNA sequencing

We used a SMART-Seq v4 Ultra Low Input RNA Kit for Sequencing (TaKaRa) for cDNA amplification according to manufacturer’s instructions. Through Columbia’s Genome Center, we prepared the library using Nextera XT DNA Library Preparation kit (Illumina) with 100-150pg starting material, then followed manufacturer’s instructions. Libraries were sequenced to a targeted depth of ∼40M 100bp paired-end reads on a NovaSeq 6000. We use RTA (Illumina) for base calling and bcl2fastq2 (version 2.19) for converting BCL to fastq format, coupled with adaptor trimming.

### RNA-seq analysis

RNA-seq reads were quantified with Kallisto^76^, using the Ensembl v96 Human:GRCh38.p12 transcriptome as a reference. Data was deposited in the Gene Expression Omnibus (GSE227571). Multidimensional scaling ^77^, principal component analysis ^78^, and hierarchical clustering^79^ were used to check for outliers. Samples were normalized by the Trimmed Mean Method^80^. Differential expression was analyzed with limma-voom with sample weighting ^81–83^ and duplicate correlation ^84^, which models the effect of 3 tissue samples per patient IgG sample. Differential expression results were analyzed with the KEGG ^85^, and Biological Process Gene Ontology ^86^, using a significance cutoff of p ≤ 0.001 and |log2 fold change| ≥0.6. iPathwayGuide ^87^ was used to perform the KEGG analysis, using the SIPA (Signal Interaction Pathway Analysis) ^88,89^ method, as well as the Gene Ontology analysis, using the High Specificity Pruning method, which is based on the elim method ^90^.

### Metabolic analysis

An extracellular flux analyzer (Seahorse XF96, Agilent) was used with the Seahorse XF Cell Mito Stress Test assay (Agilent, cat. no. 103015-100) to evaluate mitochondrial function. Prior to the assay, monolayer hiPSC-cardiomyocytes were cultured with patient IgG for 5 days. Two days prior to the assay, the cells were dissociated and replated onto an XF96 cell culture microplate (Agilent) coated with Matrigel at a cell density of 80,000 cells per XF96 well. For the remaining two days, the cells continued to be cultured in media treated with antibodies. One hour prior to the assay, the culture medium was exchanged for assay medium (XF RPMI Medium, pH 7.4 supplemented with 1 mM pyruvate, 2 mM glutamine, and 10 mM, glucose) and the culture plate was placed in a non-CO_2_ incubator. During the Mito Stress Test, electron transport chain modulators Oligomycin (1.5 µM), FCCP (0.5 µM), and Rotenone and antimycin A (0.5 µM) were injected serially into each well. The oxygen consumption rate (OCR) values were normalized to the average concentration of dsDNA present in each well of the plate, which were determined using the Quanti-iT PicoGreen dsDNA Assay Kit (Thermo Fisher Scientific cat. no. P7589). The key parameters of mitochondrial function determined by the Mito Stress Test, including basal respiration, ATP-linked respiration, proton leak, maximal respiration, spare capacity, and non-mitochondrial oxygen consumption were generated using the Seahorse XF Mito Stress Test Report Generator.

### Cardiomyocyte and cardiac fibroblasts imaging assays

hiPSC-cardiomyocytes were dissociated following differentiation and replated in Matrigel-coated 96 well plates at 200,000 cells/cm^2^. After one week, cells were cultured with patient IgG (0.5 μg/mL) for 72 hours and then analyzed. The Apotracker Green fluorescent probe (BioLegend, cat. no. 427401), MitoTracker Red CMXRos dye (Thermo Fisher cat. no. M7512), and CellROX Deep Red Reagent (Thermo Fisher cat. no. C10422) were used to quantify apoptosis, mitochondrial content, and ROS, respectively, all according to the manufacturers’ protocols. Briefly, cells were cultured with 500 nM Apotracker Green, 200 nM MitoTracker, or 5 mM CellROX reagent in culture medium for 1 hour at 37°C. Wells were washed with PBS, fixed with 4% PFA for 15 minutes at room temperature, and then washed 3 more times with PBS. Cardiac fibroblasts were cultured with patient IgG for 72 hours and immunostained as previously described in the method section to evaluate number of ki67+ cells. Primary antibody against Ki67 was used (Invitrogen, cat. no. 14-5698-82). Primary Cells were stained with DAPI and then imaged with a high throughput imaging system (BioTek Cytation 5, Agilent).

### Measurement of mitochondrial membrane potential

hiPSC-cardiomyocytes were dissociated following differentiation and replated in Matrigel-coated 96 well plates at 200,000 cells/cm^2^. After one week, cells were cultured with patient IgG (5 μg/mL) for 72 hours and then analyzed. A JC-10 mitochondrial membrane potential assay kit (Abcam cat. no. ab112134) was used according to the manufacturer’s protocol. Briefly, cells were cultured with the diluted JC-10 dye for 45 minutes at 37°C, and then the fluorescence intensity was measured on a microplate reader with 485nm/528nm and 530nm/590nm (Ex/Em) filters (BioTek Synergy HTX, Agilent). The 529nm/590nm ratio correlates with the degree of mitochondrial membrane depolarization. Incubation with 10 mM carbonyl cyanide 3-cholophenylhydrazone (Sigma-Aldrich, cat. no. C2759) for 30 minutes was used as a positive control.

### PhIP-Seq Analysis

PhIP-seq analysis and sequencing were performed according to previous publications^38,91^, with a few modifications. Sera containing approximately 100 µg of IgG (quantified using a Human IgG ELISA Quantification Set – Bethyl Laboratories) were added to each well, in duplicate, combined with 1 mL of Human PhIP-seq library^38^ diluted to approximately 2×10^5^ fold representation in phage extraction buffer (20 mM Tris-HCl, pH 8.0, 100 mM NaCl, 6 mM MgSO4). After incubation, phages were pulled down using anti-FLAG magnetic beads. IgGs that did not bind to phages were discarded and, after washing and releasing of complex phage IgG from beads, IgGs bound to phages were pulled down using anti-protein A and protein G magnetic beads. Phage DNA was extracted by incubation at 95°C for 10 min.

The DNA for multiplexed Illumina sequencing was prepared following two rounds of amplification using Q5 Hot Star Polymerase (New England Biomedical). For the first round of PCR, we used the primers IS7 (ACACTCTTTCCCTACACGACTCCAGTCAGGTGTGATGCTC) and IS8 (GTGACTGGAGTTCAGACGTGTGCTCTTCCGATCCGAGCTTATCGTCGTCATCC) and, for the second round was used primer IS4 (AATGATACGGCGACCACCGAGATCTACACTCTTTCCCTACACGACTCCAGT) and a different unique indexing primer for each sample to be multiplexed for sequencing. Equimolar amounts of all samples were pooled, gel purified and sequenced by the Harvard Medical School Biopolymers Facility using a 50-bp read cycle on an Illumina NextSeq.

### PhIP-Seq Data Analysis

Read counts were normalized by zscore and a t test was applied to compare each pair of groups (SLE versus Control, Myo-versus Myo+SD-, Myo-versus Myo+SD+ and Myo+SD-versus Myo+SD+). To compare more than two pairs, ANOVA test was applied. Peptides with pvalue lower than 0.05 were considered different for following analysis.

